# Hydrodynamic model of fish orientation in a channel Flow

**DOI:** 10.1101/2021.11.11.468193

**Authors:** Maurizio Porfiri, Peng Zhang, Sean D. Peterson

**Author notes:** **For correspondence:** (MP); (SDP).

## Abstract

For over a century, scientists have sought to understand how fish orient against an incoming flow, even without visual and flow cues. Here, we elucidate a potential hydrodynamic mechanism of rheotaxis through the study of the bidirectional coupling between fish and the surrounding fluid. By modeling a fish as a vortex dipole in an infinite channel with an imposed background flow, we establish a planar dynamical system for the cross-stream coordinate and orientation. The system dynamics captures the existence of a critical flow speed for fish to successfully orient while performing cross-stream, periodic sweeping movements. Model predictions are examined in the context of experimental observations in the literature on the rheotactic behavior of fish deprived of visual and lateral line cues. The crucial role of bidirectional hydrodynamic interactions unveiled by this model points at an overlooked limitation of existing experimental paradigms to study rheotaxis in the laboratory.

## Introduction

Swimming animals display a complex behavioral repertoire in response to flows (***Chapman et al., 2011***). Particularly fascinating is the ability of several fish species to orient and swim against an incoming flow, a behavior known as rheotaxis. While intuition may suggest that vision is necessary for fish to determine the direction of the flow, several experimental studies of midwater species swimming in a channel have documented rheotaxis in the dark above a critical flow speed (***Coombs et al., 2020***). When deprived of vision, fish lose the ability to hold station and they may perform sweeping, cross-stream movements from one side of the channel to other (***Bak-Coleman et al., 2013***; ***Bak-Coleman and Coombs, 2014***; ***Elder and Coombs, 2015***) (Fig. 1).

**Figure 1.**
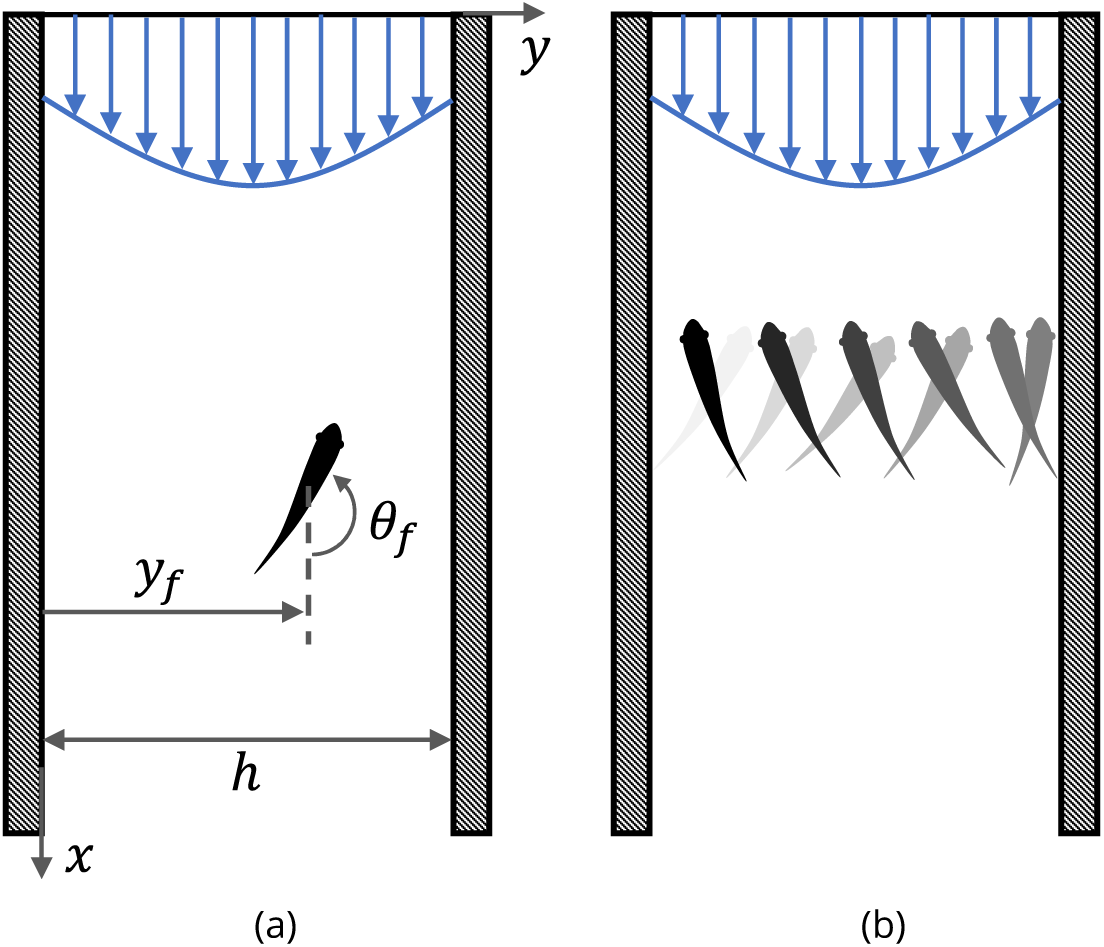
Fish rheotaxis. (a) Illustration of the problem with notation, showing a fish swimming in a background flow described by Eq. (4). (b) Schematic of the cross-stream sweeping movement of some fish species swimming without visual cues; snapshots of fish at earlier time instants are illustrated by lighter shading.

In addition to vision, fish may rely on an array of compensatory sensory modalities to navigate the flow, which utilizes tactile, proprioceptive, olfactory, electric, kinematic, and hydrodynamic signals (***Montgomery et al., 2000***; ***Von der Emde, 1999***). For example, fish could sense and actively respond to linear accelerations caused by the surrounding flow using their vestibular system (***Pavlov and Tjurjukov, 1995***). Similarly, with the help of tactile sensors on their body surface, fish could maintain their orientation against a current through momentary contacts with their surroundings (***Lyon, 1904***; ***Arnold, 1969***). Several modern studies have unveiled the critical role of the lateral line system, an array of mechanosensory receptors located on the surface of fish body (***Montgomery and Baker, 2020***), in their ability to orient against a current (***Montgomery et al., 1997***; ***Baker and Montgomery, 1999***), hinting at a hydrodynamics-based rheotactic mechanism that has not been fully elucidated. When deprived of vision, can fish rely only on lateral line feedback to perform rheotaxis? Is there a possibility for rheotaxis to be achieved through a purely passive hydrodynamic mechanism that does not need any sensing?

Through experiments on zebrafish larvae swimming in a laminar flow in a straight tube, ***Oteiza et al. (2017***) have recently unveiled an elegant hydrodynamic mechanism for fish to actively perform rheotaxis. Utilizing their mechanosensory lateral line, fish can sense the flow along different parts of their body, which is sufficient for them to deduce local velocity gradients in the flow and adjust their movements accordingly. As further elaborated upon by ***Dabiri (2017***), the insight offered by ***Oteiza et al. (2017***) is grounded in the fundamental relationship between vorticity and circulation given by the Kelvin-Stokes’ theorem, so that fish movements will be informed by local sampling of the vorticity field. While offering an elegant pathway to explain rheotaxis, the framework of ***Oteiza et al. (2017***) does not include a way for rheotaxis to be performed in the absence of information about the local vorticity field. Several experimental studies have shown that fish can perform rheotaxis even when their lateral line is partially or completed ablated, provided that the flow speed is sufficiently large (***Bak-Coleman et al., 2013***; ***Bak-Coleman and Coombs, 2014***; ***Baker and Montgomery, 1999***; ***Elder and Coombs, 2015***; ***Montgomery et al., 1997***; ***Oteiza et al., 2017***; ***Van Trump and McHenry, 2013***).

Mathematical modeling efforts seeking to clarify the mechanisms underlying rheotaxis are scant (***Oteiza et al., 2017***; ***Burbano-L and Porfiri, 2021***; ***Colvert and Kanso, 2016***; ***Chicoli et al., 2015***), despite experiments on rheotaxis dating back more than a century (***Lyon, 1904***). A common hypothesis of existing mathematical models is that the presence of the fish does not alter the flow physics with respect to the background flow, thereby neglecting interactions between the fish and the walls of the channel. For example, the model by ***Oteiza et al. (2017***) implements a random walk in a virtual flow, matching experimental measurements of the background flow in the absence of the animal through particle image velocimetry. A similar line of approach was pursued by ***Burbano-L and Porfiri (2021***) for the study of multisensory feedback control of adult zebrafish.

Thus, according to these models, the fish acts as a perfectly non-invasive sensor that probes and reacts to the local flow environment without perturbing it. There are countless examples in fluid mechanics that could question the validity of such an approximation, from coupled interactions between a fluid and a solid in vortex-induced vibrations (***Williamson and Govardhan, 2004***) to laminar boundary layer response to environmental disturbances that range from simple decay of the perturbation to bypass transition (***Saric et al., 2002***). We expect that accounting for bidirectional coupling between the fluid flow and the fish will help clarify many of the puzzling aspects of rheotaxis.

To shed light on the physics of rheotaxis, we formulate a mathematical model based on the paradigm of the finite-dipole, originally proposed by ***Tchieu et al. (2012a***). Within this paradigm, a fish is viewed as a pair of point vortices of equal and opposite strength separated by a constant distance in a two-dimensional plane. The application of the finite-dipole has bestowed important theoretical advancements in the study of hydrodynamic interactions between swimming animals (***Gazzola et al., 2016***; ***Filella et al., 2018***; ***Kanso and Tsang, 2014***; ***Kanso and Michelin, 2019***; ***Porfiri et al., 2021***), although numerical validation of the framework against full solution of Navier-Stokes equations is lacking – conducting such a validation is also part of this study. Upon validating the dipole model, we investigate the bidirectional coupling between a fish and the surrounding fluid flow in a channel. Our work contributes to the recent literature on minimal models of fish swimming (***Gazzola et al., 2014, 2015***; ***Sánchez-Rodríguez et al., 2020***) that builds on seminal work by ***Lighthill (1975***), ***Taylor (1952***), and ***Wu (1971***) to elucidate the fundamental physical underpinnings of locomotion and inform the design of engineering systems.

We focus on an ideal condition, where fish are deprived of all sensing systems, other than the lateral line that gives them access to information about the flow. Such flow information is coupled, however, to the motion of the fish itself, which acts as an invasive sensor and perturbs the background flow. Just as fish motion influences the local flow field, so too does the local flow field alter fish motion through advection. Predictions from the proposed model are compared against existing empirical observations on fish rheotaxis, compiled through a comprehensive literature review of published work since 1900. Data presented in the literature are used to offer context to the predicted dependence of rheotaxis performance on local flow characteristics, individual fish traits, and lateral line feedback.

## Results

### Model of the fluid flow

Consider a single fish swimming in an infinitely long two-dimensional channel of width *h* (Fig. 1(a)). Let one wall of the channel be at *y* = 0 and the other at *y* = *h*, with *x* pointing along the channel. The fish position at time *t* is given by 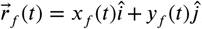, where 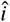 and 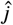 are the unit vectors in the *x* and *y* directions, respectively. The orientation of the fish with respect to the *x* axis is given by *θ*_*f*_ (*t*) (positive counter-clockwise) and its self-propulsion velocity is 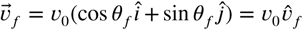, where *v*_0_ is the constant speed of the fish and 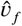 is a unit vector in the swimming direction.

The flow is modeled as a potential flow, which is a close approximation of the realistic flow field around a fish. This simple linear fluid model is intended to capture the mean flow physics, thereby averaging any turbulence contribution. The fish is modeled as a dipole, the potential field of which at some location 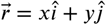 is given by

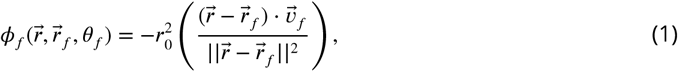

where *r*_0_ is the characteristic dipole length-scale (on the order of the amplitude of the fish tail beating), so that the circulation of each vortex is 2*πr*_0_*v*_0_. This potential field is constructed assuming a far-field view of the dipole (***Filella et al., 2018***), wherein *r*_0_ is small in comparison with the characteristic flow length scale, which is satisfied for *ρ* = *r*_0_/*h* ≪ 1. The velocity field at 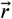 due to the dipole (fish) is 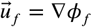.

A major contribution of the proposed model is the treatment of the fish as an invasive sensor that both reacts to and influences the background flow, thereby establishing a coupled interaction between the fish and the surrounding environment. A fish swimming in the vicinity of a wall will induce rotational flow near the boundary. In the inviscid limit, this boundary layer is infinitesimally thin and can be considered as wall-bounded vorticity (***Batchelor, 2000***). Employing the classical method of images (***Newton, 2011***), the influence of the wall-bounded vorticity on the flow field is equivalent to that of a fictitious fish (dipole) mirrored about the wall plane. For the case of a fish in a channel, this results in an infinite number of image fish (dipoles) (Fig. 2), the position vectors for which are

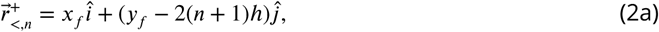

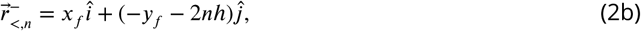

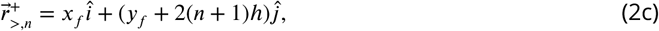

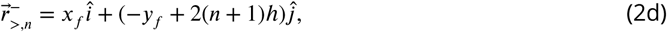

where *n* is a non-negative integer representing the *n*-th set of images. Subscripts “<“ and “>“ correspond to position vectors of the images at *y* < 0 and *y* > *h*, respectively. Likewise, superscript “±” denotes the orientation of the image dipole as ±*θ*_*f*_ ; that is, a position vector with superscript “+” indicates that the associated image has the same orientation as the fish.

**Figure 2.**
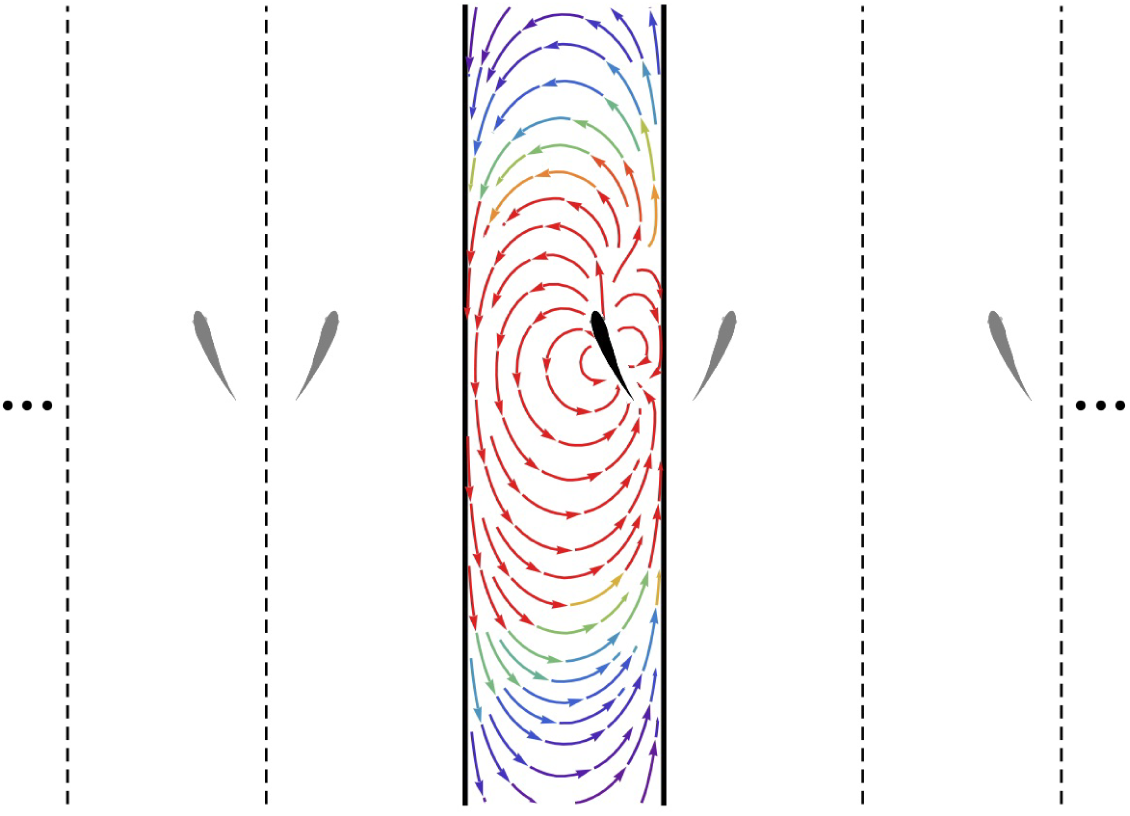
Method of images. Schematic of the fish (black) in the channel (thick lines) and the set of images (gray) needed to generate the channel. The streamlines generated by the fish in an otherwise quiescent fluid are shown in the channel colored by local velocity magnitude (red: high; blue: low). Dashed and solid lines are mirroring planes for the method of images, the pattern for which continues *ad infinitum*.

The potential function for a given image is found by replacing 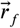 in (1) with its position vector from (2) and adjusting the sign of *θ*_*f*_ in (1) to match the superscript of its vector. The potential field at 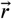 due to the image dipoles is

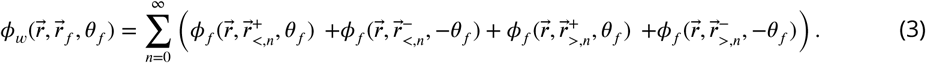

Thus, the velocity field due to the wall is computed as 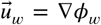, and the overall velocity field induced by the fish is 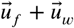. (A closed-form expression for the series in terms of trigonometric and hyperbolic functions is presented in the Supplementary Information.) Overall, the presence of the walls distorts the flow generated by the dipole, both compressing the streamlines between the fish and the walls in its proximity and creating long-range swirling patterns in the channel (Fig. 2).

The presence of a background flow in the channel is modelled by superimposing a weakly rotational flow,

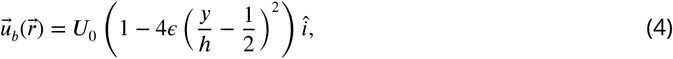

which has speed *U*_0_ at the channel centerline and *U*_0_(1 − *ϵ*) at the walls, *ϵ* being a small positive parameter. As *ϵ* → 0, a uniform (irrotational) background flow is recovered: such a flow is indistinguishable from the one in Fig. 2, provided that the observer is moving with the background flow.

For *ϵ* ≪ 1, the imposed velocity profile approximates that of a turbulent channel flow, wherein a modest degree of velocity profile curvature is present near the channel centerline. We note that this velocity profile does not satisfy the no-slip boundary condition (zero velocity on the walls), and the flow is entirely described by only two parameters (*U*_0_ and *ϵ*). For *ϵ* ≃ 1, the profile approaches that of a laminar flow with parabolic dependence on the cross-stream coordinate. The overall fluid flow in the channel is ultimately computed as 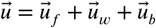.

The circulation in a region ℛ in the flow field centered at some location *y* is Γ = *∫*_ℛ_ *ω* d*A*, where 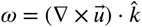 is the local fluid vorticity 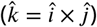. For the considered flow field, we determine

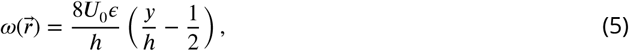

whereby the irrotational component of the flow field does not contribute to the circulation, and the circulation at a point (per unit area) is equivalent to the local vorticity.

### Numerical validation of the dipole model

Despite the success of dipole-based models in the study of fish swimming (***Tchieu et al., 2012a***; ***Gazzola et al., 2016***; ***Filella et al., 2018***; ***Porfiri et al., 2021***), their accuracy against complete Navier-Stokes simulations remains elusive. The potential flow framework in which these models are grounded neglects boundary layers and the resulting wakes that emerge from viscous effects. Quantifying the extent to which these effects influence the flow field generated by the fish is part of this study.

Specifically, computational fluid dynamics (CFD) simulations were conducted to detail the flow field around a fish during steady swimming. The simulation setup was based upon a giant danio of body length *l* = 7.3 cm in a channel of width *h* = 15.0 cm. The length of the simulation domain was *L* = 50.0 cm (∼ 6.8*l*) with the fish model placed at the channel centerline 20 cm (∼ 2.7*l*) downstream of the inlet. The body undulation of the giant danio was imposed *a priori* in the simulation, based on data from ***Najafi and Abtahi (2022***). The time-resolved flow field around the fish was quantified by solving the incompressible Navier-Stokes equations. Details on the setup of the numerical framework and convergence analysis supporting its accuracy are included in the Supplementary Information.

The mean velocity field averaged over a tail beating cycle is displayed in Fig. 3(a). The predominant flow feature observed in the mean field is a flow circulation from the head to the tail of the fish with left-right symmetry and compression of the streamlines near the channel walls. The highest velocity is found at the head of the fish with a thrust wake and recirculation region downsteam of the animal – both at a lower velocity than the anterior flow. The largest velocity in the wake is less than 20% of the peak values recorded ahead of the fish. Additional simulation results are included in the Supplementary Information. The flow field predicted by the dipole model is in good agreement with numerical simulations, as shown in Fig. 3. The dipole model is successful in capturing the circulation from the head to the tail and the compression of the streamlines near the walls. These features are expected to be the main drivers of the interaction between the fish and the channel walls, thereby supporting the value of a dipole model for a first-order analysis of the hydrodynamics of rheotaxis.

**Figure 3.**
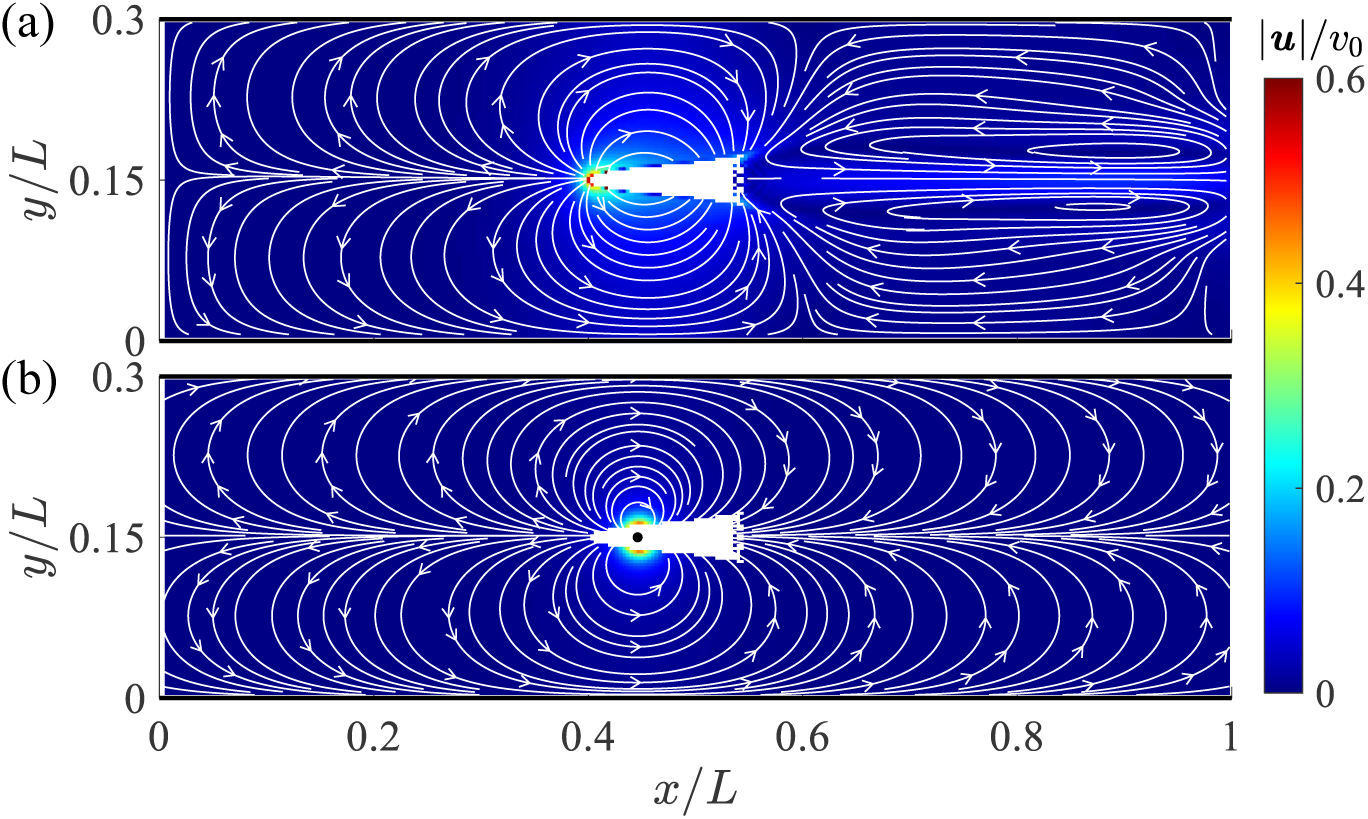
Velocity field around a swimming fish from computational fluid dynamics. (a) Mean velocity field around the steady swimming giant danio relative to the background flow. (b) Velocity field predicted by a dipole with *θ* = *π* located at 0.315*l* from the fish head along its centerline relative to the background flow. The selection of the dipole location and strength is detailed in the Supplementary Information.

### Model of fish dynamics

From knowledge of the fluid flow in the channel, we compute the advective velocity 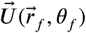 and hydrodynamic turn rate 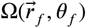 at the fish location, which encode the influence of the confining walls and background flow on the translational and rotational motion of the fish, respectively. Neglecting the inertia of the fish so that it instantaneously responds to changes in the fluid flow, we determine (***Filella et al., 2018***)

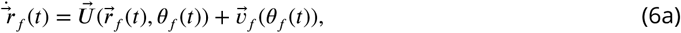

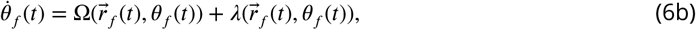

where *λ* is the feedback mechanism based on the circulation measurement through the lateral line.

The advective velocity is found by de-singularizing the total velocity field 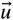 at 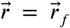, which is equivalent to calculating the sum of the velocity due to the walls and the background flow in correspondence of the fish (***Milne-Thomson, 1996***)

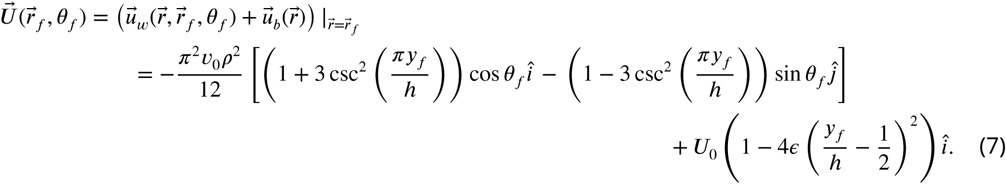

Equation (7) indicates that the walls have a retarding effect on the swimming speed of the fish that increases in magnitude the closer the fish gets to either wall of the channel. A fish swimming with orientation *θ*_*f*_ = 0 at the center of the channel, for example, will swim with velocity 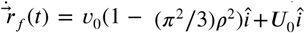. This effect should not be mistaken as traditional viscous drag, which is not included in potential flow theory; rather, it should be intended as the impact of nearby solid boundaries.

Hydrodynamic turn rate is incorporated by considering the difference in velocity experienced by the two constituent vortices comprising the dipole, namely,

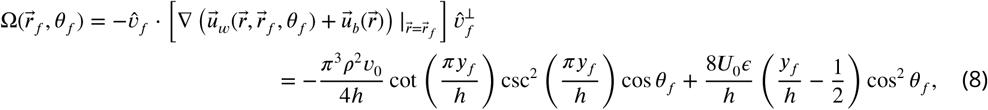

where 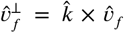; see Methods and Materials Section for the mathematical derivation. Equation (8) indicates that interaction with the walls causes the fish to turn towards the nearest wall; for example, a fish at *y*_*f*_ = 3/4*h*, will experience a turn rate due to the wall of (*π*^3^*ρ*^2^*v*_0_)/(2*h*) cos *θ*_*f*_ , such that it will be rotated counter-clockwise if swimming downstream and clockwise if swimming upstream. On the other hand, the turning direction imposed by the background flow is always positive (counter-clockwise) in the right half of the channel and negative (clockwise) in the left half, irrespective of fish orientation, so that a fish at *y*_*f*_ = 3/4*h* will always be rotated counter-clockwise. As a result, the fish may turn towards or away from a wall, depending on model parameters and orientation.

Based on experimental observations and theoretical insight (***Burbano-L and Porfiri, 2021***; ***Oteiza et al., 2017***), we hypothesize that hydrodynamic feedback, that is, lateral line measurements of the surrounding fluid that fish can employ to navigate the flow, is related to the measurement of the circulation in a region surrounding the fish. We consider a rectangular region ℛ of width *r*_0_ along the fish body length *l*. For simplicity, we assume a linear feedback mechanism, 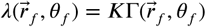, where we made evident that circulation is computed about the fish location and *K* is a non-negative feedback gain. Assuming that the fish size is smaller than the characteristic length scale of the flow, we linearize the vorticity along the fish in (5) as 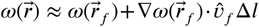. By computing the integral from Δ*l* = −*l*/2 to *l*/2, we obtain

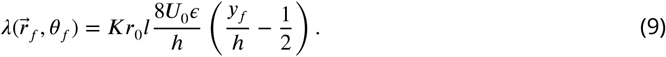

Compared to established practice for modeling fish behavior in response to visual stimuli (***Gautrais et al., 2009***; ***Calovi et al., 2014***; ***Zienkiewicz et al., 2015***; ***Couzin et al., 2005***), the proposed model introduces nonlinear dynamics arising from the bidirectional coupling between the motion of the fish and the flow physics in its surroundings. We note that the employed feedback in (9) neglects additional potential sensing mechanisms, including vision (***Lyon, 1904***), acceleration sensing through the vestibular system (***Pavlov and Tjurjukov, 1995***), and pressure sensing through sensory afferents in the fins (***Hardy et al., 2016***), which might enhance the ability of fish to navigate the flow.

### Analysis of the planar dynamical system

Given that the right hand side of equation set (6) is independent of the streamwise position of the fish, the equations for the cross-streamwise motion and the swimming direction can be separately studied, leading to an elegant nonlinear planar dynamical system. We center the cross-stream coordinate about the center of the channel and non-dimensionalize it with respect to *h*, introducing *ξ* = *y*_*f*_ /*h* − 1/2. The governing equations become

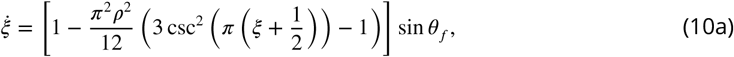

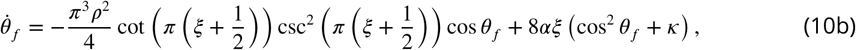

where we non-dimensionalized by the time needed for the fish to traverse the channel in the absence of a background flow, that is, *h*/*v*_0_, and introduced *α* = *U*_0_*ϵ*/*v*_0_ and *κ* = *Kr*_0_*l* (see Methods and Materials Section for estimation of these parameters from experimental observations).

In search of the equilibria of the dynamical system, we note that swimming downstream or upstream (*θ*_*f*_ = 0 and *π*, respectively) solves (10a) for any choice of the cross-stream coordinate, the value of which is determined from the solution of (10b) for the corresponding orientation *θ*_*f*_. In the case of downstream swimming, the only solution of the resulting transcendental equation is *ξ* = 0. For upstream swimming, depending on the value of the parameter *β* = (*α*(1 + *κ*))/*ρ*^2^, we have one or three solutions: if *β* < *β*^*^ = *π*^4^/32, the only solution is *ξ* = 0, otherwise, in addition to *ξ* = 0, there are two solutions symmetrically located with respect to the centerline that approach the walls as *β* → ∞ (Fig. 4(a), see Methods and Materials Section for mathematical derivations).

**Figure 4.**
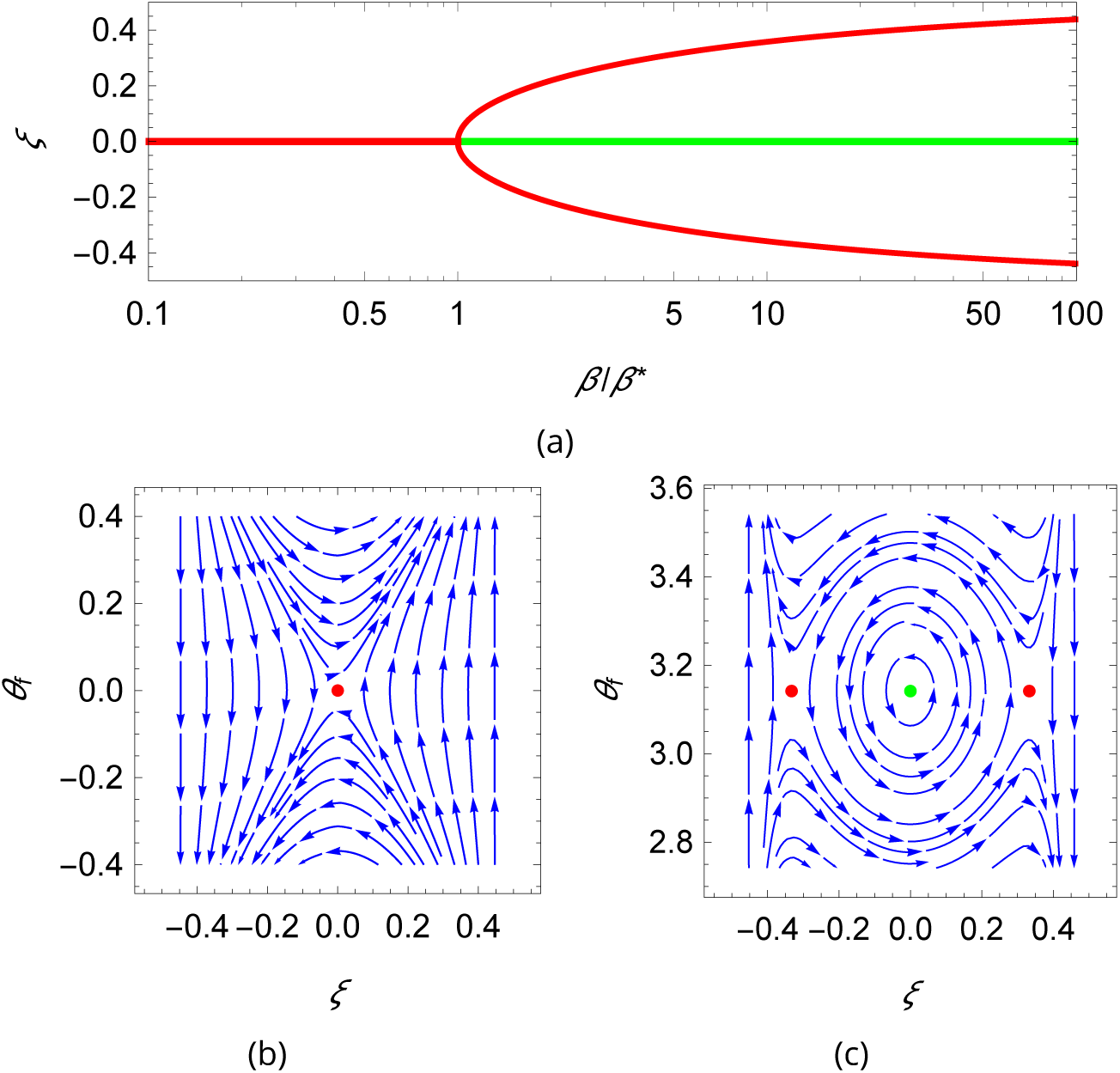
Qualitative dynamics of equation set (10). (a) Cross-stream equilibria for upstream swimming as a function of *β*. (b,c) Phase plot for downstream and upstream swimming in the case *α* = 0.1, *ρ* = 0.01, and *κ* = 1, so that *β* = 20. In all panels, red refers to unstable equilibria and green to stable equilibria.

Local stability of these equilibria is determined by studying the eigenvalues of the state matrix of the corresponding linearized dynamics. For all the considered dynamics, the trace of the state matrix is zero, so that the equilibria can be saddle points (unstable) or neutral centers (stable), if the determinant is negative or positive, respectively (***Bakker, 1991***) (see Methods and Materials Section for mathematical derivations). In the case of downstream swimming, the determinant is always negative, such that the equilibrium (*θ*_*f*_ = 0, *ξ* = 0) is a saddle point (Fig. 4(b)). For upstream swimming, the equilibrium (*θ*_*f*_ = *π, ξ* = 0) is stable if *β* > *β*^*^, leading to periodic oscillations similar to experimental observations (***Bak-Coleman et al., 2013***; ***Bak-Coleman and Coombs, 2014***; ***Elder and Coombs, 2015***) (Fig. 1(b)); the other two equilibria located away from the centerline are always unstable (Figs. 4(b,c)). Oscillations about the centerline during rheotaxis have a radian frequency 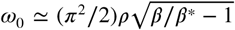, such that the frequency increases with the square root of *β* and is zero at *β*^*^ (see Methods and Materials Section for the mathematical derivation).

## Discussion

There is overwhelming evidence that fish can negotiate complex flow environments by responding to even small flow perturbations (***Liao, 2007***). However, seldom are these perturbations included in mathematical models of fish behavior, which largely rely on vision cues (***Gautrais et al., 2009***; ***Calovi et al., 2014***; ***Zienkiewicz et al., 2015***; ***Couzin et al., 2005***). In this paper, we proposed a hydrodynamic model for the bidirectional coupling between fish swimming and fluid flow in the absence of any sensory input but lateral line feedback – encapsulated by a simple linear feedback mechanism. The model reduces to a nonlinear planar dynamical system for the cross-stream coordinate and orientation, of the kind that are featured in nonlinear dynamics textbooks for their elegance, analytical tractability, and broad physical interest (***Sastry, 2013***).

The planar system anticipates several of the surprising features of rheotaxis. In particular, this study provides some potential answers to the question raised by ***Coombs et al. (2020***): “…what role, if any, do passive (e.g. wind vane) mechanisms play in rheotaxis and how are these influenced by fish factors (e.g. body shape) and flow dynamics?” Through the mathematical analysis of the model, we uncovered an equilibrium at the channel centerline for upstream swimming whose stability is controlled by a single non-dimensional parameter that summarizes flow speed, lateral line feedback, flow gradient, channel width, and fish size. Above a critical value of this parameter, the model predicts that rheotaxis is stable and fish will begin periodic cross-stream sweeping movements whose amplitude can be as large as the channel width. Interestingly, the model anticipates rheotaxis even without sensory feedback, through only passive hydrodynamic mechanisms.

Our mathematical proof of the existence of a nontrivial threshold for *β* above which upstream swimming becomes stable finds partial support in experimental observations on a number of species in the absence of visual cues (see Supplementary Information, where we have performed a bibliographical survey on experimental studies about rheotaxis). Several of these experiments have indicated the existence of a threshold in the flow speed or flow gradient above which fish successfully perform rheotaxis. Importantly, we predict that the presence of channel walls is necessary for the emergence of such a threshold, since for *ρ* → 0, *β* → ∞, thereby automatically guaranteeing the stability of upstream swimming. Based on our estimation of *α* and *ρ* from available data, *β* can be as small as 10^−1^ and exceed 10^2^, thereby encompassing the critical value *β*^*^ ≃ 3 (see Methods and Materials Section for estimation of model parameters). We should exercise care in drawing comparisons with experiments, which only control for visual feedback, in contrast with the model where we block all sensory modalities except of the lateral line. As reviewed by ***Coombs et al. (2020***), water-motion cues can also be accessible to tactile or other cutaneous senses, beyond the lateral line that is included in our model. In addition, body-motion cues are not limited to visual senses, whereby they can be accessed by tactile and vestibular senses. Hence, a one-to-one comparison between experiments and theory is presently not possible.

The model predicts the emergence of rheotaxis in the absence of any sensory information. Setting *κ* = 0 in our model eliminates hydrodynamic feedback, yet, the fish is able to perform rheotaxis at sufficiently large flow speeds and steep flow gradients. This finding would support the possibility of a completely passive mechanism for rheotaxis. To date, there is no experiment on live animals that can be used to support this claim, owing to the necessity to eliminate all sensory modalities without compromising fish ability to swim. In practice, this may be unfeasible to do. As discussed in ***Coombs et al. (2020***), existing approaches for disabling senses suffer from potential pitfalls, including: i) unintended effects on the overall fish behavior, which are likely to occur in an effort to block at once vestibular, tactile, and lateral line senses, and ii) difficulty in guaranteeing complete blockage of a sensory modality, which, like the lateral line, can be distributed throughout the whole body.

A potential line of approach to explore the possibility of a complete passive form of rheotaxis is through experiments with robotic fish (***Zhang et al., 2016***; ***Wang et al., 2020***; ***Duraisamy et al., 2019***), mimicking locomotory patterns of live animals and allowing to precisely control sensory input. In this vein, we foresee experiments with robotic fish in a complete open-loop operation that does not utilize any sensory input. The robotic fish developed in our previous study (***Kopman and Porfiri, 2013***; ***Kopman et al., 2014***) could offer a versatile platform to conduct such an experiment. Such a robot is actuated by a built-in step motor to undertake a periodic tail beating with a predetermined frequency. All its electronics is encased in the frontal section of the robot, so that its size and shape can be readily adjusted though rapid prototyping.

Although free swimming experiments on the robotic fish would be ideal, practicality may suggest to constraint the streamwise and vertical location of the robot while allowing cross-stream motions and heading changes. Such a setup shall also include a load cell to measure the drag on the robot, providing an independent measure to set the tail beat frequency – similar to CFD simulations (see Supplementary Information). Also, we would recommend measuring energy expenditure by the robotic fish to gain insight into the hydrodynamic costs and benefits of rheotaxis. Experimental parameters encapsulated in *β*, including inlet flow speed, channel width, and robot length can be all controlled, and the flow curvature at the centerline can be measured through velocimetry techniques. In the experiments, one shall track the motion of the robotic fish to score conditions in which stable rheotaxis is observed and extract other salient information, such as the frequency of cross-stream sweeping, if present.

The model predicts that increasing *κ* broadens the stable region, leading to more robust rheotaxis, which is in qualitative agreement with experimental observations – including experiments on animals with intact versus compromised lateral lines (see Supplementary Information). The model prediction on the influence of the environment on rheotaxis, including the flow gradient and flow channel size, also parallels the literature on rheotaxis (see Supplementary Information). For example, observations by ***Oteiza et al. (2017***) suggest that increasing the flow gradient *ϵ* enhances hydrodynamic feedback in zebrafish, resulting in improved rheotaxis.

Our analysis also indicates that wider channels should promote rheotaxis by lowering the critical speed above which swimming against the flow becomes a stable equilibrium. This mathematical finding is indirectly supported by experimental observations (see Supplementary Information), and bears relevance in the design of experimental protocols for the study of rheotaxis. Confining the subject in a narrow channel will promote bidirectional hydrodynamic interactions with the walls, so that small movements of the animal will reverberate into sizeable changes in the flow physics that will mask the gradient of the background flow. Similarly, in partial alignment with experimental observations, the model predicts a lower threshold for longer fish, owing to a magnification of the hydrodynamic feedback received by a longer body (see Supplementary Information). Again, we warn care in drawing comparisons due to the presence of other senses in real experiments, which are not modelled in our work.

Finally, the model anticipates the onset of periodic cross-stream sweeping, which has been studied in some experiments on fish swimming in channels without vision (***Coombs et al., 2020***). While there is not conclusive experimental evidence regarding the dependence of the frequency of oscillations on flow conditions, the model is in qualitative agreement with experiments by ***Elder and Coombs (2015***), showing a sublinear dependence on the flow speed. Therein, it is shown that the radian frequency has a weak positive tendency with respect to the flow speed for Mexican tetra swimming with or without cues from the lateral line. Above 2 cm s^−1^, the animals can successfully perform rheotaxis and display sweeping oscillations at about three cycles per minute and increase to about four cycles per minute at 12 cm s^−1^. These correspond to a radian frequency on the order of 0.1 rad s^−1^, which is similar to what we would predict for *β* ranging from 10^0^ to 10^1^ and *ρ* of the order of 10^−1^ (recall that the time is scaled with respect to time required by the animal to traverse the channel from wall to wall in the absence of a background flow). We acknowledge that the current model does not describe contact and impact with the walls of the channel, which could be important in further detailing the onset of cross-sweeping motions that could involve stick-and-slip at the bottom of the channel (***Van Trump and McHenry, 2013***).

Just as other minimal models of fish swimming have helped resolve open questions on scaling laws (***Gazzola et al., 2014***), gait (***Gazzola et al., 2015***), and drag (***Sánchez-Rodríguez et al., 2020***), the proposed effort addresses some of the baffling aspects of rheotaxis through a transparent and intuitive treatment of bidirectional hydrodynamic interactions between fish and their surroundings. The crucial role of these bidirectional interactions hints that active manipulation of their surroundings by fish offers them a pathway to overcome sensory deprivation and sustain stable rheotaxis.

The proposed model is not free of limitations, which should be addressed in future research. The current model neglects the elasticity and inertia of the fish, which might reduce the accuracy in the prediction of rheotaxis, especially transient phenomena. Future research should refine the dipole paradigm toward a dynamic, unsteady model that accounts for added mass effects and distributed elasticity, similar to those used in the study of swimming robots (***Sfakiotakis et al., 1999***; ***Colgate and Lynch, 2004***). The model could also be expanded to account for additional sensory modalities, such as vision, vestibular system, and tactile sensors on the fish body surface. We argue that pursuing any of these extensions shall require detailed experimental data, beyond what the literature can currently offer. Experiments should also be needed to refine the linear hydrodynamic feedback mechanism that we hypothesized for the lateral line; in this vein, future experiments could be designed to parametrically vary the flow speed and quantify the activity level of lateral line nerve fibers through neurophysiological recordings (***Mogdans, 2019***). Beyond the inclusion and refinement of individual sensory modalities, we envision research toward the incorporation of a multisensory framework, as the one introduced by ***Coombs et al. (2020***). Such a framework identifies several motorsensory integration sites in the central nervous system that could contribute to rheotaxis, thereby calling for modeling efforts at the interface of neuroscience and fluid mechanics.

Despite its limitations, the proposed minimalistic model is successful in anticipating some of the puzzling aspects of rheotaxis and points at the possibility of attaining rheotaxis in a purely passive manner, without any sensory input. Most importantly, the model brings forward a potential methodological oversight of laboratory practice in the study of rheotaxis, caused by bidirectional hydrodynamic interactions between the swimming fish and the fluid flow. To date, there is no gold standard for the selection of the size of the swimming domain, which is ultimately chosen on the basis of practical considerations, such as facilitating behavioral scoring and creating a laminar background flow. The model demonstrates that the width of the channel has a modulatory effect on the threshold speed for rheotaxis and the cross-stream swimming frequency, which challenges the comparison of different experimental studies and confounds the precise quantification of the role of individual sensory modalities on rheotaxis. Overall, our effort warrants reconsidering the behavioral phenotype of rheotaxis, by viewing fish as an invasive sensor that modifies the encompassing flow and hydrodynamically responds to it.

## Methods and Materials

### Derivation of the turn rate equation for the fish dynamics

The expression for the turn rate in equation (8) is obtained from the original finite-dipole model by ***Tchieu et al. (2012b***), in the limit of small distances between the vortices in the pair (*r*_0_ → 0).

Specifically, equation (2.11) from ***Tchieu et al. (2012b***), adapted to the case of a single dipole reads

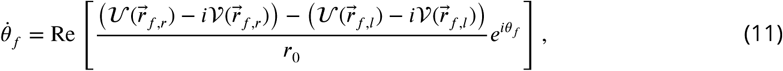

where subscript *l* and *r* refer to the left and right vortices forming the pair and 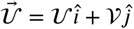 is the advective velocity field acting on the dipole. The advective field consists of the interactions with the walls and the background flow, so that 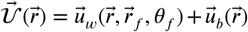; in the case of ***Tchieu et al. (2012b***), such a field encompasses the velocity field induced by any other dipole in the plane. Left and right vortices are defined so that 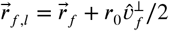 and 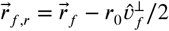, which yields 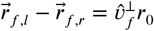.

By carrying out the complex algebra in (11), we determine

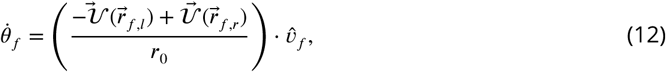

which supports the intuition that the dipole will turn counter-clockwise if the right vortex would experience a stronger velocity along the swimming direction. Upon linearizing the term in parenthesis in the neighborhood of 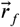, this expression becomes

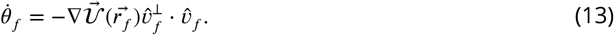

The chosen approach is consistent from the standpoint of vortex dynamics, by which each vortex in the pair advects in response to local fluid velocity. In this vein, the fish is interpreted as a bluff body, which rotates according to a difference in the drag experienced by its left and right sides. Such a difference is amplified by the pectoral fins, which enhance the effect of any left-to-right asymmetry in the surrounding fluid flow. In the literature, this description is termed T-dipole, in opposition to the so-called A-dipole that introduces two fiducial points along the direction of motion of the dipole that govern its turning (***Kanso and Tsang, 2014***). Whether one representation is superior to the other in terms of accuracy is yet to be clarified; our choice of using a T-dipole is based on its theoretical consistency and intuition on the underlying flow physics. Potential avenues for resolution include detailed CFD simulations of free swimming fish or experiments with robotic fish.

### Determination of the equilibria of the planar dynamical system

By setting *θ*_*f*_ = 0 or *θ*_*f*_ = *π* in equation set (10), we determine that *ξ* should be equal to some constant, which is a root of the following transcendental equation:

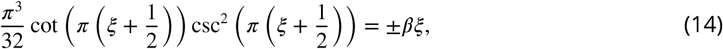

where the positive sign corresponds to *θ*_*f*_ = 0 and the negative sign to *θ*_*f*_ = *π*. Here, *β* = *α*(1 + *κ*)/*ρ*^2^ as introduced from the main text.

As shown in Fig. 5, for *θ*_*f*_ = 0, there is only one root of the equation (*ξ* = 0; see the intersection between the solid red line and the solid black curve), while up to three roots can rise for *θ*_*f*_ = *π* depending on the value of *β*. For *β* smaller than a critical value *β*^*^, only *ξ* = 0 is a solution (see the intersection between the solid blue line and the solid black curve), while for *β* > *β*^*^ two additional solutions, symmetrically located with respect to the origin emerge (see the intersections between the dashed blue line and the solid black curve). The critical value *β*^*^ is identified by matching the slope of the black curve at *ξ* = 0, so that *β*^*^ = *π*^4^/32. Notably, the two solutions symmetrically located with respect to the centerline approach the walls as *β* → ∞.

**Figure 5.**
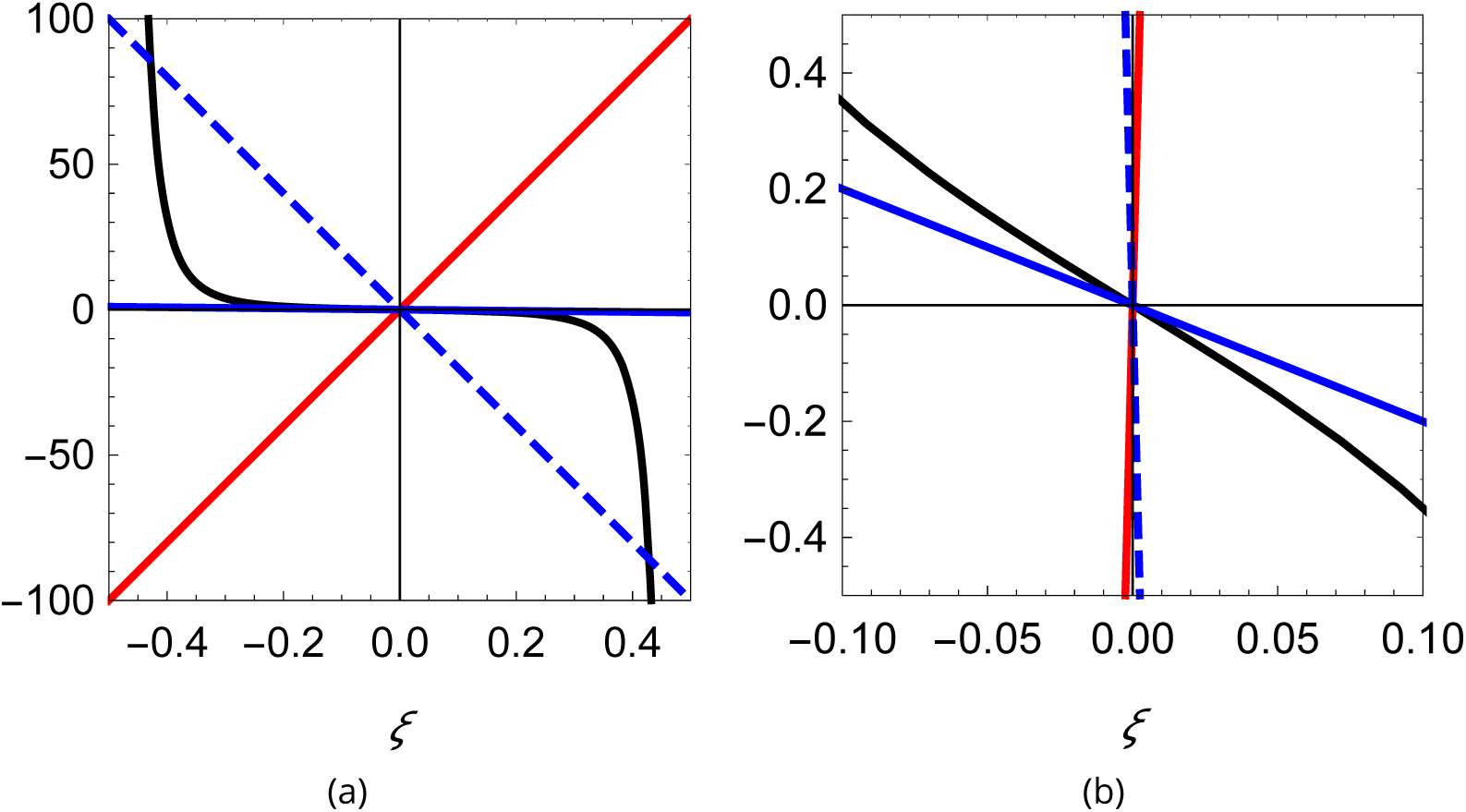
Visual illustration of the process of determining the roots of (14). (a) Plot of the function 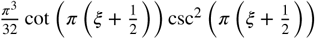 (black), superimposed with three lines of different slope: 200 (red), −200 (dashed blue), and −2 (solid blue). (b) Zoomed-in view of the curves in (a) showing that the blue line can only intersect the black curve at the origin.

### Local stability analysis of the planar dynamical system

To examine the local stability of the equilibria of the planar dynamical system, we linearize equation set (10). The state matrix of the linearized dynamics, *A*, describes the local behavior of the nonlinear system when perturbed in the vicinity of the equilibrium, that is,

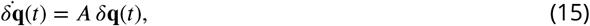

where *δ***q** = [*δξ, δθ*_*f*_]^T^ is the variation about the equilibrium. The eigenvalues of the *A* are indicative of local stability about each equilibrium.

For *θ*_*f*_ = 0 and *ξ* = 0, the state matrix is given by

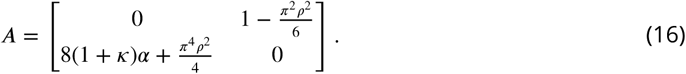

Given that the trace of the matrix is zero (tr *A* = 0), the analysis of the stability of the equilibrium resorts to ensuring the sign of the determinant to be positive (det *A* > 0). Specifically, if the determinant is positive, the eigenvalues are imaginary and the equilibrium is a neutral center (stable, although not asymptotically stable), otherwise one of the eigenvalues is positive and the equilibrium is a saddle point (unstable) (***Bakker, 1991***). Hence, stability requires that

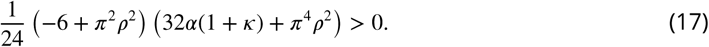

Since the first factor is always negative (*ρ* ≪ 1) and the second is positive, the inequality is never fulfilled and the equilibrium is a saddle point (unstable) (Fig. 4(a,b)).

For *θ*_*f*_ = *π* and *ξ* = 0, the state matrix is given by

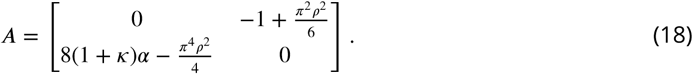

Similar to the previous case, stability requires that det *A* > 0, that is,

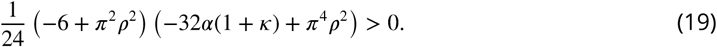

Due to the sign change in the first summand appearing in the second factor with respect to the previous case, stability becomes possible. Specifically, the equilibrium is a neutral center (stable) for *β* > *β*^*^ = *π*^4^/32, which is also the necessary condition for the existence of the two equilibria symmetrically located with respect to the channel centerline (Fig. 4(a,c)).

When *β* > *β*^*^, we register the presence of two more equilibria at ±*ξ* ≠ 0. The state matrix takes the form

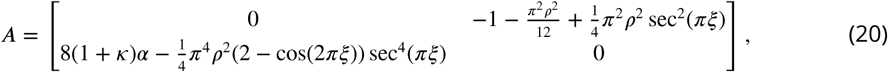

Also in this case, stability requires that det *A* > 0, that is,

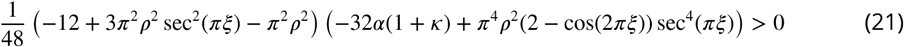

Once again, for *ρ* ≪ 1, we can assume that the first factor in parenthesis is negative. (This assumption is grounded upon (14), which yields that (*ξ* ± 1/2) = 𝒪 (*ρ*^2/3^); since cos(*πξ*)^2^ = 𝒪 ((*ξ* ± 1/2)^2^), we have that *ρ*^2^ sec^2^(*πξ*) → 0 as *ρ* → 0.) Hence, we obtain

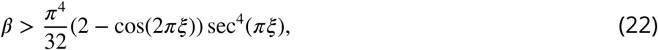

which is not satisfied for any choice of *β* > *β*^*^. Thus, the two equilibria away from the channel centerline, close to the walls are always saddle points (unstable) (Figs. 4(a,c)).

We comment that the local stability analysis requires only knowledge of the curvature of the flow field at the centerline of the channel. Hence, should one contemplate alternative profiles for the background flow, linear stability results shall not change. Higher-order parameterizations for the flow profile will result into nonlinear dependencies on *ξ* that do not affect the linear analysis. Likewise, while we considered a linear feedback mechanism to integrate lateral information via a simple gain, one may explore nonlinear relationships between *λ* and Γ. The linear stability analysis shall not change, whereby these nonlinear forms will result into dependencies on higher powers of *ξ*.

### Frequency of cross-stream sweeping

The linearized planar system about the stable focus in (18) is equivalent to a classical second-order system in terms of the cross-stream coordinate, similar to a mass-spring model. Hence, the radian resonance frequency of the system is

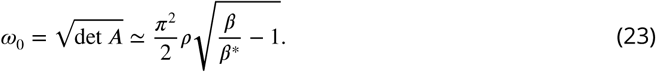

where the last approximation holds for *ρ* ≪ 1. Equation (23) shows that, close to the threshold, the frequency of oscillations is small and it increases with *β* and *ρ*.

### Estimation of model parameters

In a typical experimental setup on rheotaxis, the width of the channel, *h*, is on the order of three to ten times the body length of the animal, *l*. For example, experiments from ***Elder and Coombs (2015***) on Mexican tetras of *l* = 8.3 cm were conducted in a channel with *h* = 25 cm. Similarly, in the experiments on adult zebrafish from ***Burbano-L and Porfiri (2021***), *l* = 3.6 cm and *h* = 13.8 cm, and in the experiments on zebrafish larvae from ***Oteiza et al. (2017***), *l* = 4.2 mm (inferred from the animals’ age) and *h* = 1.27 – 4.76 cm. The distance between the vortices simulating a fish, *r*_0_, should be on the order of a tail beat, which has a typical value of 0.2*l* (***Gazzola et al., 2014***). As a result, it is tenable to assume that *ρ*^2^ is between 10^−4^ and 10^−2^.

A safe estimation of the velocity of the animal in the absence of the background flow, *v*_0_, would be on the order of few body lengths per second (***Gazzola et al., 2014***). The speed used for the background flow across experiments, *U*_0_, tend to be of the same order as the magnitude of *v*_0_, leaning toward values close to one body length per second (***Coombs et al., 2020***). For instance, data on zebrafish from ***Burbano-L and Porfiri (2021***) suggest *v*_0_ = 5.7 cm s^−1^ and *U*_0_ = 3.2 cm s^−1^. The estimation of the non-dimensional parameter *ϵ* associated with the shear in the flow is more difficult, since data on the velocity profiles are seldom reported. That being said, for channel flow of sufficiently high Reynolds number, the velocity profile in the channel is expected to be blunt, approximating a uniform flow profile near the channel center (***White, 1974***). Thus, it is tenable to treat *ϵ* as a small parameter, between 10^−2^ and 10^−1^. For flow of low Reynolds number (***Oteiza et al., 2017***) (Re < 100), the velocity gradient in the channel has been observed to be large, corresponding to *ϵ* values in the range of 10^−1^ and 1. By combining these estimations, we propose that *α* ranges between 0 and 1.

An estimation of *κ* is difficult to offer, whereby feedback from the lateral line has only been included in few studies (***Oteiza et al., 2017***; ***Burbano-L and Porfiri, 2021***; ***Colvert and Kanso, 2016***; ***Chicoli et al., 2015***). Using the data-driven model from ***Burbano-L and Porfiri (2021***), it is tenable to assume values on the order of 10^1^ for individuals showing high rheotactic performance. This gain can also be estimated by comparing the threshold speeds of fish, *U*_*c*_, with and without the lateral line, through 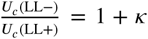, according to (S4) in the Supplementary Information. The significant increase in the threshold speed following lateral line ablation in ***Baker and Montgomery (1999***) indicates that *κ* ∈ [2, 7], while the indistinguishable threshold speed between LL+ and LL− fish in a few other studies (***Bak-Coleman and Coombs, 2014***; ***Elder and Coombs, 2015***; ***Van Trump and McHenry, 2013***) may suggest that *κ* ∼ 0. In Table 1, we summarize the model parameters identified from data in the experimental studies detailed in the Supplementary Information.

**Table 1.** Estimation of model parameters from data in the literature.

| Reference | $\rho$ | $\epsilon$ | $\alpha$ | $\kappa$ | $\beta$ |
| --- | --- | --- | --- | --- | --- |
| Bak-Coleman et al. (2013) | $\sim 0.05$ | $[10^{-2}, 10^{-1}]$ | $[0, 0.17]$ | — | — |
| Bak-Coleman and Coombs (2014) | $\sim 0.04$ | $[10^{-2}, 10^{-1}]$ | $[0, 0.16]$ | $\sim 0$ | $[0, 100]$ |
| Baker and Montgomery (1999) and Montgomery et al. (1997) | $\sim 0.1$ | $[10^{-2}, 10^{-1}]$ | $^\dagger[0, 0.32]$ | $[2, 7]$ | $[0, 256]$ |
| Elder and Coombs (2015) | $\sim 0.066$ | $[10^{-2}, 10^{-1}]$ | $^\dagger[0, 0.24]$ | $\sim 0$ | $[0, 55]$ |
| Kulpa et al. (2015) | $\sim 0.04$ | $\sim 1$ near center of jet | $\sim 1.3$ | — | — |
| Oteiza et al. (2017) | $[0.018, 0.066]$ | $[0.20, 0.82]$ | — | — | — |
| Peimani et al. (2017) | $\sim 0.044$ | $\sim 1$ | — | — | — |
| Suli et al. (2012) | $\sim 0.018$ | $[0.1, 1]$ | — | — | — |
| Van Trump and McHenry (2013) | $[0.055, 0.127]$ | $[10^{-2}, 10^{-1}]$ | $^\dagger[0, 0.32]$ | $\sim 0$ | $[0, 106]$ |

$^\dagger$ LL+ cavefish swimming speed $v_0 \sim 5 \text{ cm/s}$ in zero background flow in Bak-Coleman and Coombs (2014) is used to estimate $\alpha$

## Acknowledgments

The authors acknowledge financial support from the National Science Foundation under Grant No. CMMI-1901697. The authors acknowledge Alain Boldini and Simone Macrì for useful discussions.

## Author Contributions

M.P. and S.D.P. conceived the study. M.P. and S.D.P. developed the theoretical model and performed data analysis. P.Z. conducted the literature review and performed the model validation.

M.P. and S.D.P. wrote a first draft of the manuscript, which was consolidated in its present form by all the authors.

## Data and materials availability

The authors declare that the data supporting the findings of this study are available within the paper. The Mathematica notebook used to derive the governing equations, study the planar dynamical, and generate associated figures, together with the CFD data discussed in the text, are also available at https://github.com/dynamicalsystemslaboratory/Rheotaxis.

