## Supplementary Information for "Hydrodynamic model of fish orientation in a channel Flow"

[1]Department of Biomedical Engineering, New York University Tandon School of Engineering; 370 Jay Street, 11201 Brooklyn (NY) - USA; [2]Department of Mechanical and Aerospace Engineering, New York University Tandon School of Engineering; 370 Jay Street, 11201 Brooklyn (NY) - USA; [3]Center for Urban Science and Progress, New York University Tandon School of Engineering; 370 Jay Street, 11201 Brooklyn (NY) - USA; [4]Mechanical and Mechatronics Engineering Department, University of Waterloo; 200 University Avenue West, Waterloo ON N2L 3G1 - Canada

### Complete expression for the velocity field caused by image dipoles

The velocity field $\vec{u}_f = u_f \hat{\imath} + v_f \hat{\jmath}$ at $\vec{r}$ induced by the single dipole at $\vec{r}_f$, given by the potential function in equation (1), is

$$u_f(\vec{r},\vec{r}_f,\theta_f) = r_0^2 v_0 \left( \frac{((x-x_f)^2-(y-y_f)^2)\cos\theta_f + 2(x-x_f)(y-y_f)\sin\theta_f}{((x-x_f)^2+(y-y_f)^2)^2} \right), \tag{S1a}$$

$$v_f(\vec{r},\vec{r}_f,\theta_f) = r_0^2 v_0 \left( \frac{-((x-x_f)^2-(y-y_f)^2)\sin\theta_f + 2(x-x_f)(y-y_f)\cos\theta_f}{((x-x_f)^2+(y-y_f)^2)^2} \right). \tag{S1b}$$

The potential function describing the image vortex system for a dipole in a channel presented in equation (3) can be simplified using `Mathematica`, yielding

$$\phi_w(\vec{r},\vec{r}_f,\theta_f) = \frac{r_0^2 v_0}{4}\left[ 4\frac{(x-x_f)\cos\theta_f + (y-y_f)\sin\theta_f}{(x-x_f)^2+(y-y_f)^2} - \frac{\pi e^{-i\theta_f}}{h}\left(e^{2i\theta_f}\left(\coth(\pi A)+\coth(\pi B^*)\right)+\coth(\pi A^*)+\coth(\pi B)\right)\right], \tag{S2}$$

where $A = ((x-x_f)+i(y-y_f))/(2h)$, $B = ((x-x_f)+i(y+y_f))/(2h)$, $i=\sqrt{-1}$, and a superscript * indicates complex conjugate. The velocity field at $\vec{r}$ due to the walls, $\vec{u}_w = u_w\hat{\imath} + v_w\hat{\jmath}$, is

$$u_w = \frac{r_0^2 v_0}{4}\left[ \frac{\pi^2 e^{-i\theta_f}}{2h^2}\left(e^{2i\theta_f}(\operatorname{csch}^2\pi A + \operatorname{csch}^2\pi B^*) + \operatorname{csch}^2\pi A^* + \operatorname{csch}^2\pi B\right) + \frac{4\cos\theta_f}{(x-x_f)^2+(y-y_f)^2} - \frac{8(x-x_f)((x-x_f)\cos\theta_f+(y-y_f)\sin\theta_f)}{((x-x_f)^2+(y-y_f)^2)^2}\right], \tag{S3a}$$

$$v_w = \frac{r_0^2 v_0}{4}\left[ \frac{i\pi^2 e^{-i\theta_f}}{2h^2}\left(e^{2i\theta_f}(\operatorname{csch}^2\pi A - \operatorname{csch}^2\pi B^*) - \operatorname{csch}^2\pi A^* + \operatorname{csch}^2\pi B\right) + \frac{4\sin\theta_f}{(x-x_f)^2+(y-y_f)^2} - \frac{8(y-y_f)((x-x_f)\cos\theta_f+(y-y_f)\sin\theta_f)}{((x-x_f)^2+(y-y_f)^2)^2}\right]. \tag{S3b}$$

Superimposing the velocity fields from the dipole and its images and setting $y=0$ (or $y=h$) yields $v_f + v_w = 0$, thereby confirming that the walls of the channel are streamlines.

### Computational fluid dynamics

#### Framework

To quantify the flow around the swimming fish, we solved the incompressible Navier-Stokes equations numerically in the commercial software COMSOL Multiphysics (version 5.6). We focused on the steady swimming of a giant danio exhibiting a carangiform swimming pattern, consisting of large body undulations in the posterior of the animal and minimal lateral movement in the anterior portion. The lateral movement of a giant danio was mathematically described through a local coordinate system, $x'$-$y'$, such that the undeformed fish aligns its centerline along the $x'$-axis with head at $x' = 0$ and tail at $x' = l$ (see Fig. 1(a)). The lateral movement were described as a traveling wave from the head to the tail as (***Najafi and Abtahi, 2022***)

$$y' = \left[c_1 \left(x' - 0.3l\right) + c_2 \left(x' - 0.3l\right)^2\right] \sin\left[k_L \left(x' - 0.3l\right) - 2\pi f t\right], \tag{S4}$$

where $c_1$ and $c_2$ are two constants that describe the shape of the undulation envelope, $k_L$ is the wavenumber, and $f$ is the tail beating frequency. The values of $l$, $c_1$, $c_2$, and $k_L$ are taken from ***Najafi and Abtahi*** (***2022***), and the value of $f$ is taken from ***Zhang et al.*** (***2019***); these parameters are summarized in Table S1. For the chosen tail beat frequency, the oscillatory Reynolds number ***Gazzola et al.*** (***2014***) is $\mathrm{Sw} = \frac{2\pi C f l}{\nu} = 8,100$, where $C$ is the tail beat amplitude, given as $C = 0.7 \times c_1 l + 0.49 \times c_2 l^2$, and $\nu$ is the kinematic viscosity of water at room temperature ($\nu = 1.0 \times 10^{-6}\,\mathrm{m^2 s^{-1}}$).

**Table S1.** Parameters employed in (S4) to describe the locomotory pattern of a giant danio.

| Parameters | $l$ | $c_1$ | $c_2$ | $k_L$ | $f$ |
|---|---|---|---|---|---|
| Values | 7.3 cm | 0.004 | $-2.33\,\mathrm{m^{-1}}$ | $78.5\,\mathrm{m^{-1}}$ | 3 Hz |

The undeformed giant danio body shape was approximated as a NACA0013 airfoil, which has a maximum thickness-to-chord length matching that of the animal in ***Zhang et al.*** (***2019***). The movement of the fish body was imposed in the numerical simulations using a moving boundary. No-slip boundary conditions was set on the fish body, whereas the channel walls satisfied only the no-penetration boundary condition, that is, they were treated as slip walls. In the simulations, we set a uniform flow with speed $U_0$ at the inlet, imposed a zero pressure boundary condition at the outlet, and fixed the axial location of the fish at the channel centerline.

The fluid domain was discretized using triangular elements, as illustrated in Fig. 1(a), and was allowed to deform in time to accommodate the body undulations. A grid convergence study was performed to ensure the solution was independent of the mesh. As shown in Fig. 1(b), a total number of $76\,\mathrm{k}$ elements were sufficient to guarantee the mesh independence of the simulations, such that implementing a higher number of $122\,\mathrm{k}$ elements introduced negligible variation in the mean drag prediction on the fish. As a result, all simulations were conducted using $76\,\mathrm{k}$ elements. A total of $10$ tail beating cycles were simulated in each simulation, which was sufficiently long for the flow in the channel to fully develop. All simulations were conducted on $12$ Intel Xeon Platinum $8268$ CPUs with a base frequency of $2.90\,\mathrm{GHz}$ and a total of $48\,\mathrm{GB}$ memory. A typical simulation required approximately $6\,\mathrm{h}$ computational time to complete.

To represent the realistic swimming condition, the imposed inlet flow speed, $U_0$, should match the fish swimming speed, $v_0$, which depends on the body undulations described by (S4). To identify the value of $v_0$, we conducted a series of preliminary simulations by varying $U_0$ until the time-averaged total drag on the fish, $\bar{F}_\mathrm{D}$ was zero. That is, the drag experienced by the fish exactly balanced the thrust generated by the body undulations. The resulting swimming speed was determined to be $v_0 = 19.74\,\mathrm{cm/s}$. The resulting Reynolds number based upon channel width is $\mathrm{Re} = \frac{hU_0}{\nu} = 29,600$.

#### Results

The instantaneous velocity fields in the vicinity of the giant danio relative to the background flow are presented at five time instants during half of a tail beating cycle in Fig. S2. In comparison with

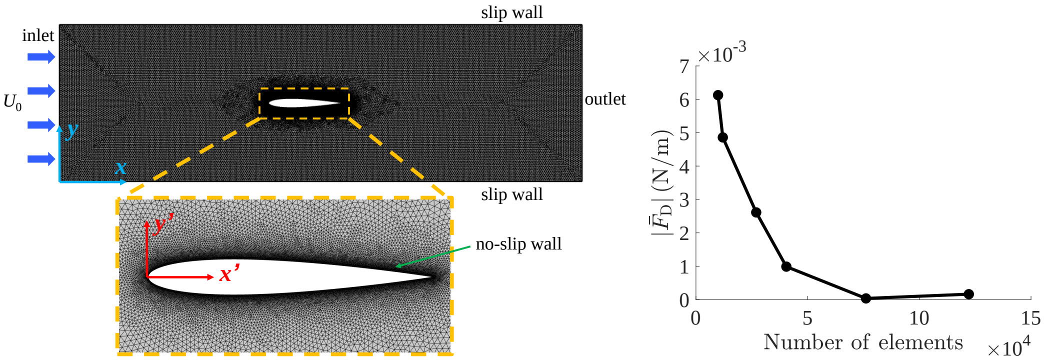


(a) (b)

**Figure S1.** Details about the implementation of the computational fluid dynamics simulations. (a) Mesh implemented in the simulations, with definitions of coordinate systems and a zoomed-in view of the refined mesh around the fish. (b) Mesh convergence analysis, showing the mean drag as a function of number of elements in the simulation.

---

the mean flow shown in Fig. 3, we observe distortions of the streamlines, with a left-right asymmetry caused by the body undulations. We also identify a series of alternating vortices generated through tail beating that form the wake. The magnitude of the wake flow decays as it is advected downstream.

CFD results are also useful to quantify the thickness of the boundary layer along the animal and offer support in favor of the proposed inviscid model for the study of the interactions between a fish and the channel walls. The thickness of the boundary layer can be quantified from the flow velocity component along the $x$-direction, $u_{\mathrm{t}}$, in the presence of the background flow. As shown in Fig. S3, the value of $u_{\mathrm{t}}$ is zero at the body surface due to the no-slip boundary condition. A large gradient is observed within a thin layer at the boundary, in which $u_{\mathrm{t}}$ rapidly increases to $v_0$. The boundary layer thickness can be estimated by identifying the location at which $u_{\mathrm{t}}$ reaches $99\%$ of its asymptotic value away from the fish body. During a tail beating cycle, the boundary layer thickness ranged between $2.8\%$ and $15.7\%$ of the fish body length, thereby supporting the feasibility of neglecting the influence of the viscous boundary layers as a first-order approximation.

### Comparison with the dipole model

The velocity field predicted by the dipole model is validated against the mean velocity quantified through CFD (see Fig. 3(a)). Consistent with the fish location and heading direction, we set $y_f = h/2$ and $\theta = \pi$ for the dipole. The axial location, $x_f$, and the characteristic length scale, $r_0$, of the dipole are treated as fitting parameters. The velocity field associated with the dipole model in Fig. 3(b) corresponds to a set of fitting parameters that minimizes the discrepancy between the model prediction and the simulated simulation. Denoting the model-predicted velocity field as $\vec{u}^{\mathrm{dipole}}$ and the simulated one as $\vec{u}^{\mathrm{CFD}}$, the discrepancy between them is quantified through

$$E = \sqrt{\frac{\int_D \left|\vec{u}^{\mathrm{dipole}} - \vec{u}^{\mathrm{CFD}}\right|^2 \, \mathrm{d}A}{\int_D \mathrm{d}A}}, \tag{S5}$$

where $D$ is the computational domain. The optimal values of the fitting parameters that minimize $E$ are determined through an exhaustive search of the parameter space, leading to $(x_f - x_s) = 0.315l$ and $r_0 = 0.062l$, where $x_s$ is the axial location of the fish head (the origin of the $x'$-$y'$ coordinate system).

### Bibliographical survey

We surveyed over three hundred publications cited by ***Arnold*** (***1974***) and ***Coombs et al.*** (***2020***) – two review papers on rheotaxis, with the former focusing on early investigations from 1900s to 1970s,

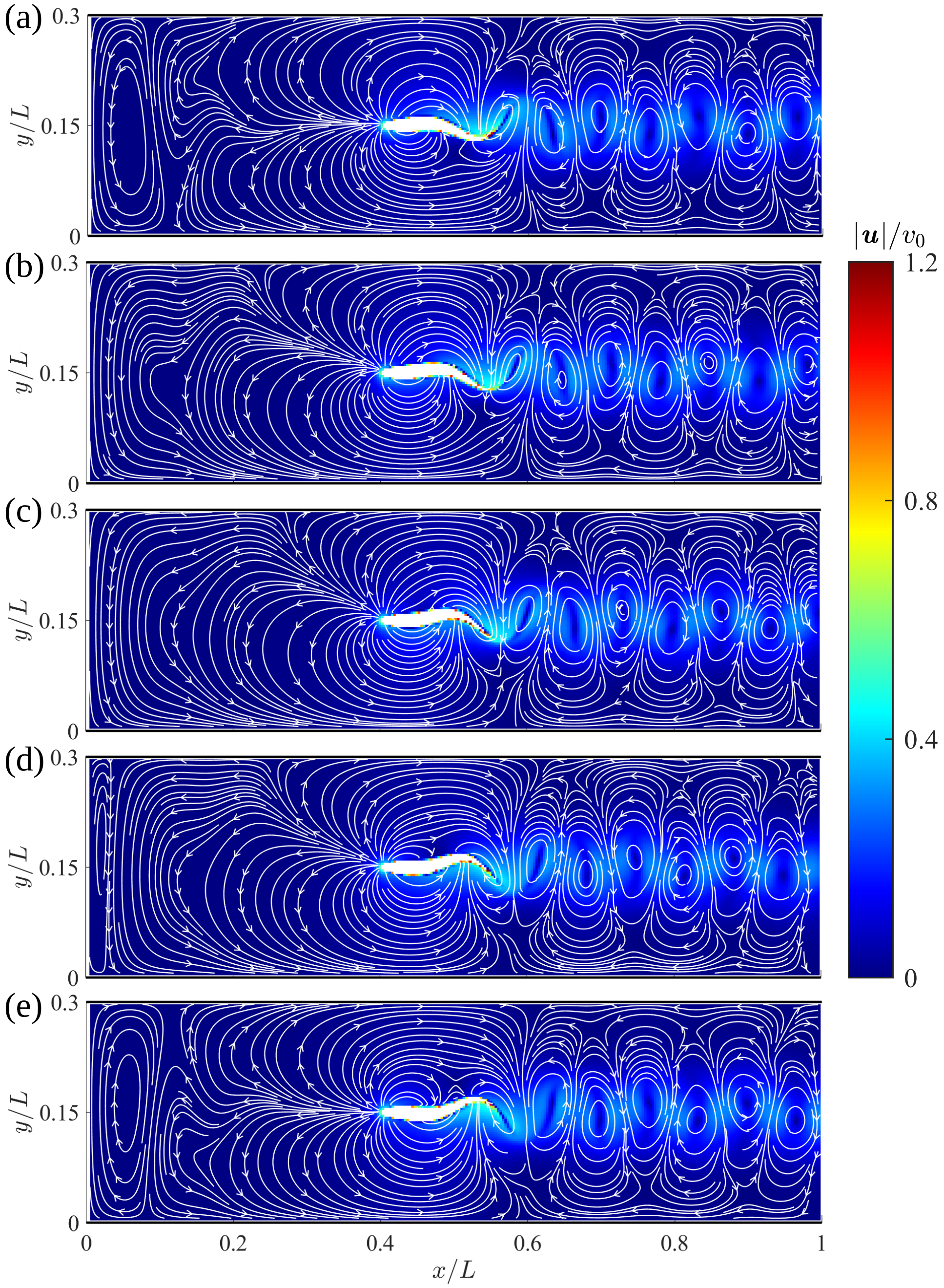


**Figure S2.** Instantaneous velocity fields around a swimming fish relative to the background flow from computational fluid dynamics. (a) – (e) correspond to $t = 0$, $T/8$, $T/4$, $3T/8$, and $T/2$, respectively, where $T$ is the period of a tail beat. White curves are streamlines with arrows indicating flow directions.

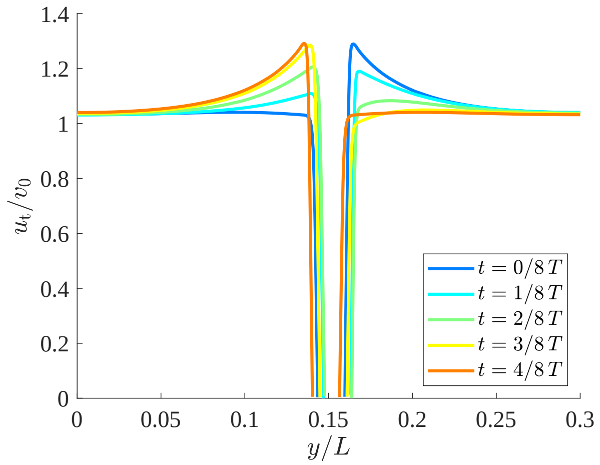


**Figure S3.** Analysis of the boundary layer thickness along a swimming fish. $x$-component of the flow velocity, $u_t$, extracted across the half-length of the fish body. The values of $u_t$ are measured in a coordinate system moving at the speed of $v_0$ along the fish swimming direction.

and the latter highlighting more recent works conducted between 1970s and 2020. Publications were selected through the following inclusion and exclusion criteria.

### Criteria

*Inclusion criteria:* We selected studies where: i) the subject animals were fish; ii) fish demonstrated rheotactic behavior; iii) no unsteady flow events were present in the swimming domain, such as the wake structure of obstacles; iv) the sensory cues available to fish could be identified with some confidence; v) fish behavior was not influenced by social interactions; vi) fish swam without visual cues; and vii) the publication was written in English. Within criterion iii), we focused on experiments with steady flows where the flow gradient is consistent over time, thus excluding swimming in random flow events. Criterion vi) was introduced to direct our search toward the effects of hydrodynamic cues and lateral line sensing, which limited our search to experiments using blind fish or experiments in the dark. We acknowledge that this condition is not reflective of the model hypotheses, which, in fact, block any sense except for the lateral lines (such as vestibular and tactile senses).

*Exclusion criteria:* Among studies identified through the selection criteria, we excluded experiments on pleuronectiform flatfishes, which swim on their side and generate propulsive undulations in a vertical plane (***Webb, 2002***). This locomotory pattern differs fundamentally from the current model, derived on the assumption the fish align their bodies vertically and undulate on a horizontal plane, which is the swimming strategy of the majority of fishes.

### Dataset

Table S2 presents data extracted from the selected studies, including the fish species, size of the swimming domain, flow conditions, sensory cues available to the fish, and the measured rheotaxis threshold speed. Swimming domains with rectangular cross-sections are defined by their length ($L$), width ($h$), and depth ($W$), while swimming domains with circular cross-sections by their length and diameter ($D$). Flow conditions are quantified through the flow speed and flow gradient. If information about the flow gradient was not available in a study, we qualitatively estimated its value through the Reynolds number of the flow, defined based on the width (diameter) of the channel and the background flow speed as $\mathrm{Re} = \frac{hU_0}{\nu}$ ($\mathrm{Re} = \frac{DU_0}{\nu}$). For a sufficiently high $\mathrm{Re}$, the flow gradient near the center of the channel is expected to be low.

**Table S2.** Relevant publications on fish rheotaxis in the absence of visual cues, identified through literature review.

| Reference | Fish | | Swimming domain | Flow properties | | [†]Sensory cues | Rheotaxis threshold speed |
|---|---|---|---|---|---|---|---|
| | Species | Length | | Flow speed | Flow gradient | | |
| *Bak-Coleman et al.* (*2013*) | Giant danio (*Devario aequipinnatus*) | 6.0 – 7.3 cm | Flow tank of $25 \times 25 \times 25$ cm ($L \times h \times W$) | 0, 3, and 7 cm/s | Re ∼ 7500 at LL+ threshold speed; flow gradient expected to be small near center of tank | LL+/LL– | ≤ 3 cm/s |
| *Bak-Coleman and Coombs* (*2014*) | blind cavefish (*Astyanax mexicanus*) | 4.2 – 5.0 cm | Flow tank of $25 \times 25 \times 25$ cm ($L \times h \times W$) | 0, 1, 2, 3, 4, 7, and 8 cm/s | Re ∼ 2000 at LL+ threshold speed; flow gradient expected to be small near center of tank | LL+/LL–; fish made transient contacts with substrate | LL+: 0.90 cm/s; LL–: 0.54 cm/s |
| [*]*Baker and Montgomery* (*1999*) | blind cavefish (*Astyanax fasciatus*) | 4 – 7 cm | Flow tank of $51 \times 9 \times 20$ cm ($L \times h \times W$) | 0, 2, 3, 5, 9, 16 cm/s | Re ∼ 2000 at LL+ threshold speed; flow gradient expected to be small near center of tank | LL+/LL–; tactile senses | LL+: 2–3 cm/s; LL–: 9–16 cm/s |
| *Elder and Coombs* (*2015*) | Mexican tetras (*Astyanax mexicanus*) | 8.3 cm | Flow tank of $25 \times 25 \times 25$ cm ($L \times h \times W$) | 0, 1, 2, 4, 7, and 12 cm/s | Re ∼ 5000 at threshold speed; flow gradient expected to be small near center of tank | LL+/LL– | ∼ 2 cm/s for LL+ and LL– |
| *Kulpa et al.* (*2015*) | blind cavefish (*Astyanax mexicanus*) | 4.4 – 5.3 cm | Flow tank of $25 \times 25 \times 10$ cm ($L \times h \times W$) | Maximum speed of 8 cm/s | Jet flow across center of tank; flow gradient expected to be large | LL+/LL– | ≤ 8 cm/s |
| [‡]*Lyon* (*1904*) | blind Fundulus | unspecified | Trough with unspecified dimensions; tideway leading to pond | "not too strong current" in trough and current with "more or less eddy and irregularity" in tideway | Flow gradient expected to be small | LL+; some fish gained tactile senses | Not measured; rheotaxis elicited only by tactile cues |
| [‡]*Lyon* (*1904*) | blind Fundulus | unspecified | Trough with unspecified dimensions | flow "gushing rather violently" | Jet flow; flow gradient expected to be large | LL+ | Not measured; rheotaxis elicited by flow |
| [*]*Montgomery et al.* (*1997*) | blind cavefish (*Astyanax fasciatus*) | 4 – 7 cm | [§]Flow tank of $51 \times 9 \times 20$ cm ($L \times h \times W$) | 0, 2, 3, 5, 9, and 16 cm/s | Re ∼ 2000 at LL+ threshold speed; flow gradient expected to be small near center of tank | LL+/LL–; tactile senses | LL+: 2–3 cm/s; LL–: 9–16 cm/s |

**Table S2.** Relevant publications on fish rheotaxis in the absence of visual cues, identified through literature review.

| Reference | Fish | | Swimming domain | Flow properties | | †Sensory cues | Rheotaxis threshold speed |
|---|---|---|---|---|---|---|---|
| | Species | Length | | Flow speed | Flow gradient | | |
| *Oteiza et al.* (*2017*) | zebrafish (*Danio rerio*) larva 5–7 days post fertilization (dpf) | unspecified | 13 cm-long circular tube with diameter 1.27–4.76 cm | 0.2–0.8 cm/s | Low to high flow gradients identified through particle image velocimetry | LL+/LL– | LL+: rheotaxis observed as low as 0.2 cm/s |
| *Peimani et al.* (*2017*) | zebrafish (*Danio rerio*) larva 5–7 dpf | estimated $\sim 0.35$ cm | Flow channel of $63.3 \times 1.6 \times 0.55$ mm ($L \times h \times W$) | 0.95–3.8 cm/s | $\mathrm{Re} \sim 10$ at threshold speed; flow gradient expected to be large | LL+ | 0.95 cm/s |
| *Suli et al.* (*2012*) | zebrafish (*Danio rerio*) larva 5 dpf | $\sim 0.33$ cm | Flume of $110 \times 3.7 \times 2.8$ cm ($L \times h \times W$) | 0.075, 0.15, 0.2 cm/s | $\mathrm{Re} < 75$; flow gradient expected to be large | LL+/LL– | Not quantified |
| *Van Trump and McHenry* (*2013*) | blind Mexican cavefish (*Astyanax fasciatus*) | 3 – 7 cm | Cylindrical channel of $150 \times 11$ cm ($L \times D$) | 0, 1, 2, 4, 6, 8, 10, 13, 16 cm/s | $\mathrm{Re} > 2000$ at threshold speed; flow gradient expected to be small near center of tank | LL+/LL– | 2–4 cm/s |

† LL+: lateral line enabled; LL–: lateral line disabled

* Data are extracted from the same set of experiments

‡ Two experiments are considered from the same paper

§ Data are from *Baker and Montgomery* (*1999*)

### Comparison against model predictions

The studies identified in Table S2 are utilized to offer some context to the proposed theoretical framework with respect to the rheotaxis stability threshold. We first express the stability threshold $\beta = \beta^*$ in dimensional form in terms of the rheotaxis threshold speed

$$U_c = \frac{\pi^4 r_0^2 v_0}{32h^2(1 + Kr_0 l)\epsilon}, \tag{S6}$$

such that $U_0 > U_c$ corresponds to the stable condition $\beta > \beta^*$, and vice versa. From most studies, the values of $U_c$ can be identified and its confidence level can be inferred (see Methods and Materials Section).

As evidenced in (S6), a series of parameters could influence the rheotaxis threshold speed, including the lateral line feedback, flow gradient, swimming domain size, and fish body length. Specifically, (S6) predicts that increasing the lateral line feedback, flow gradient, and/or width of swimming channel promotes rheotaxis at lower flow speeds, whereas increasing fish size will require higher flow speeds to elicit rheotactic behavior. The effects of these parameters are validated independently in Table S3, where we include experimental evidence garnered within each study and, when possible, carry out a comparison, across them. We compare each of these empirical observations to model predictions, and assess if they support the model, contradict the model, or are inconclusive. An observation is considered supportive of (contradictive to) our model if the measured $U_c$ exhibits with statistical significance the same (opposite) dependence on a certain parameter. Data that lack statistical significance are considered inconclusive.

The confidence intervals of the measured $U_c$ values were estimated to determine if $U_c$ were significantly different across studies. When the mean and standard error of the mean (s.e.m.) of $U_c$ were provided, we estimated its 95% confidence interval as $(\mathrm{mean} - 1.96\,\mathrm{s.e.m.}, \mathrm{mean} + 1.96\,\mathrm{s.e.m.})$. If the confidence intervals of two $U_c$ values did not overlap, we considered them significantly different. For instance, in ***Bak-Coleman and Coombs*** (***2014***) and ***Elder and Coombs*** (***2015***), the confidence intervals of $U_c$ were determined to be $(0.63, 1.17)\,\mathrm{cm/s}$ and $(1.27, 2.64)\,\mathrm{cm/s}$, respectively, and thus the $U_c$ values were considered significantly different. In several studies, such as ***Baker and Montgomery*** (***1999***) and ***Van Trump and McHenry*** (***2013***), the threshold speeds were only estimated as intervals, where fish swimming below a lower bound did not perform rheotaxis, while they exhibited rheotaxis above an upper bound. We treated this speed interval as the confidence interval for $U_c$ in our statistical analysis.

*Effect of lateral line feedback:* Several studies provide some support in favor of the prediction of our model of the beneficial role of lateral line feedback, showing a significant reduction in rheotactic performance when the lateral line is compromised (***Kulpa et al., 2015***; ***Oteiza et al., 2017***; ***Suli et al., 2012***), see Table S3. In these studies, fish locomotion was measured in steady background flows, so that a fish holding station would experience minimal linear acceleration and marginally engage the vestibular system. Throughout these studies, fish were not observed to make contact with the swim channel, indicating that tactile senses played a negligible role in rheotaxis.

*Effect of flow gradient:* We identified two studies (***Lyon, 1904***; ***Oteiza et al., 2017***) that could back the predicted effect of the flow velocity gradient on rheotaxis, as summarized in Table S3. In both studies, fish locomotion was recorded in flows with varying velocity gradients. In qualitative agreement with the proposed model, the rheotaxis performance of zebrafish larvae significantly improved with increasing gradient magnitudes (***Oteiza et al., 2017***). Similar observations were obtained by Lyon on blind Fundulus (***Lyon, 1904***), where in a flow with a small gradient, fish performed rheotaxis only when tactile cues were available, while in a jet flow with a large flow gradient, rheotaxis could be elicited solely by the flow. Although qualitatively in line with our predictions, we conservatively considered this study as inconclusive due to a lack of quantitative data for statistical tests.

*Effect of size of swimming domain:* To elucidate the role of the swimming domain size on rheotaxis threshold speed, we conducted cross-study comparisons as shown in Table S3. As evidenced through comparisons between two experiments on zebrafish larvae (***Oteiza et al., 2017***; ***Peimani et al., 2017***) in swim channels of drastically different sizes, rheotaxis was elicited at a higher threshold speed in a smaller flow channel, which supports our model prediction. Our model is also qualitatively supported by comparisons between ***Bak-Coleman and Coombs*** (***2014***) and ***Baker and Montgomery*** (***1999***), or ***Bak-Coleman and Coombs*** (***2014***) and ***Van Trump and McHenry*** (***2013***), where experiments on blind cavefish of comparable body sizes uncovered higher threshold speeds in smaller flow channels. In the experiments of ***Bak-Coleman and Coombs*** (***2014***), blind cavefish were observed to receive transient tactile senses while swimming, which could have contributed to its lower rheotaxis threshold. As a result, experimental data on blind cavefish were conservatively deemed to be inconclusive.

*Effect of fish size:* The relationship between the threshold speed and fish body size is less straightforward, as the body size not only determines the value of $l$, but also influences $r_0$, which is on the order of the fish tail beat amplitude. We assumed $r_0 = 0.2l$, which is a typical tail-beat-amplitude-to-body-length ratio (***Gazzola et al., 2014***). For fish with functional lateral lines that produce positive feedback, $K > 0$, we obtain $U_c \sim 1 - \frac{1}{1+0.2Kl^2}$. For fish with disabled lateral line, $K = 0$, we find $U_c \sim l^2$. In both cases, the model predicts that $U_c$ is larger for fish with larger body length. Some evidence can be garnered by contrasting a pair of studies by ***Bak-Coleman and Coombs*** (***2014***) and ***Elder and Coombs*** (***2015***), where experiments were conducted on fish of the same species (*Astyanax mexicanus*) in swim tunnels of the same size, and tested in flows at a similar range of speeds. The high flow speeds in both studies suggest that the flow gradients in these experiments were small. We assume that the lateral line feedback were equivalent in both studies, as the subjects were conspecific. Although the tactile cues present in the experiments by ***Bak-Coleman and Coombs*** (***2014***) hinder our ability to reach a definitive conclusion on the effect of fish body size, the higher threshold speed observed in larger fish qualitatively supports our model prediction. The paucity of data for validation of the effect of fish size is a result of a lack of studies with matching experimental conditions, including dimensions of the flow facilities, flow conditions, and functionality of the lateral line.

In summary, we identified a total of five sets of experiments in support of our model, and nine sets of studies that offer inconclusive evidence. None of the data contradicted predictions from the proposed model.

**Table S3.** Results of the bibliographical research on fish rheotaxis in the absence of visual cues, used to validate the proposed model.

| Reference | Fish species | [†]Evidence | Comparison with model | |
|---|---|---|---|---|
| | | | Supportive | Inconclusive |
| **Within studies** | | | | |
| *Effect of lateral line* | | | | |
| ***Bak-Coleman et al.*** (***2013***) | Giant danio (*Devario aequipinnatus*) | No significant difference in fish heading angle against current was detected between LL+ and LL− | | × |
| ***Bak-Coleman and Coombs*** (***2014***) | blind cavefish (*Astyanax mexicanus*) | Rheotaxis threshold speed was slightly (but not significantly) lower in LL− condition | | × |
| ***Baker and Montgomery*** (***1999***) and ***Montgomery et al.*** (***1997***) | blind cavefish (*Astyanax fasciatus*) | Rheotaxis threshold speed was significantly higher in LL− condition; fish received intermittent tactile senses | | × |
| ***Elder and Coombs*** (***2015***) | Mexican tetras (*Astyanax mexicanus*) | No significant influence of LL condition was detected on rheotactic performance | | × |
| ***Kulpa et al.*** (***2015***) | blind cavefish (*Astyanax mexicanus*) | Significantly higher rheotaxis index in LL+ fish than LL− fish in jet stream | × | |
| ***Oteiza et al.*** (***2017***) | zebrafish (*Danio rerio*) larva 5–7 dpf | Posterior lateral line ablation or chemical neuromast ablation severely reduced rheotaxis | × | |

**Table S3.** Results of the bibliographical research on fish rheotaxis in the absence of visual cues, used to validate the proposed model.

| Reference | Fish species | †Evidence | Comparison with model | |
|---|---|---|---|---|
| | | | Supportive | Inconclusive |
| ***Suli et al.*** (***2012***) | zebrafish (*Danio rerio*) larva 5 dpf | LL hair cell damage led to a significant decrease in rheotaxis; regeneration of LL hair cells restored rheotaxis | × | |
| ***Van Trump and McHenry*** (***2013***) | blind Mexican cavefish (*Astyanax fasciatus*) | In LL+ and LL−, fish exhibited statistically indistinguishable rheotaxis behavior | | × |
| *Effect of flow gradient* | | | | |
| ***Lyon*** (***1904***) | blind Fundulus | In a flow with small gradient, rheotaxis was elicited only when fish received tactile cues; in jet flow with large gradient, rheotaxis was elicited by flow without tactile cues. Lack of data on statistical significance | | × |
| ***Oteiza et al.*** (***2017***) | zebrafish (*Danio rerio*) larva 5–7 dpf | Rheotaxis of fish improved with increasing gradient magnitudes | × | |
| **Across studies** | | | | |
| *Effect of channel width* | | | | |
| ***Bak-Coleman and Coombs*** (***2014***); ***Baker and Montgomery*** (***1999***) | blind cavefish (*Astyanax mexicanus*); blind cavefish (*Astyanax fasciatus*) | Significantly different threshold speed for LL+ fish: $0.90 \pm 0.137\,\mathrm{cm/s}$ (mean ± s.e.m.) in $25\,\mathrm{cm}$ wide tunnel; between $2\,\mathrm{cm/s}$ and $3\,\mathrm{cm/s}$ in $9\,\mathrm{cm}$ wide tunnel. Tactile cues available to fish in ***Bak-Coleman and Coombs*** (***2014***) | | × |
| ***Bak-Coleman and Coombs*** (***2014***); ***Van Trump and McHenry*** (***2013***) | blind cavefish (*Astyanax mexicanus*); blind cavefish (*Astyanax fasciatus*) | Significantly different threshold speed for LL+ fish: $0.90 \pm 0.137\,\mathrm{cm/s}$ (mean ± s.e.m.) in $25\,\mathrm{cm}$ wide tunnel; between $2\,\mathrm{cm/s}$ and $4\,\mathrm{cm/s}$ in $\sim 11\,\mathrm{cm}$ diameter tunnel. Tactile cues available to fish in ***Bak-Coleman and Coombs*** (***2014***) | | × |
| ***Oteiza et al.*** (***2017***); ***Peimani et al.*** (***2017***) | zebrafish (*Danio rerio*) larva 5–7 dpf | Onset of rheotaxis in LL+ fish observed at flow speed $0.95\,\mathrm{cm/s}$ in $1.6\,\mathrm{mm}$ wide tunnel; rheotaxis observed in LL+ fish at flow speed $0.2\,\mathrm{cm/s}$ in $2.22\,\mathrm{cm}$ diameter tunnel | × | |
| *Effect of body length* | | | | |
| ***Bak-Coleman and Coombs*** (***2014***); ***Elder and Coombs*** (***2015***) | blind cavefish (*Astyanax mexicanus*); Mexican tetras (*Astyanax mexicanus*) | Significantly different threshold speed for LL+ fish: $0.90 \pm 0.137\,\mathrm{cm/s}$ (mean ± s.e.m.) for 4.2–$5.0\,\mathrm{cm}$ long fish; $1.96 \pm 0.350\,\mathrm{cm/s}$ (mean ± s.e.m.) for $8.3\,\mathrm{cm}$ long fish. Tactile cues available to fish in ***Bak-Coleman and Coombs*** (***2014***) | | × |
| **Total** | | | 5 | 9 |

† LL+: lateral line enabled; LL−: lateral line disabled

Most experiments used in our comparison listed in Table S2 were conducted in the past 25 years, and only two studies date back to before 1970. This disparity is attributed to an evolution of the methodologies for the study of rheotaxis over time. Among the earlier efforts, a large portion relied on observations of fish behavior in the field (***Arnold, 1974***). Although these studies minimized the introduction of external stimuli stemming from human presence and unfamiliar environments that could alter the behavior of fish in the wild, a lack of flexibility in the design of controlled experiments in the field, together with an insufficient measurement resolution, has led to only a limited number of works that could distinguish the impact of one sensory cue from another. As a result, a large number of earlier efforts do not meet our inclusion criteria.

Likely, the interest in fish rheotaxis was recently reignited owning to the advancements in technologies that could selectively deactivate specific fish sensory organs, thereby allowing for the targeted investigation of the role of each sensory cue in rheotaxis. For instance, pharmacological methods that could disable the lateral line led to studies (***Montgomery et al., 1997***; ***Baker and Montgomery, 1999***) challenging the long-standing perception that the lateral line could not mediate rheotaxis. In addition, the development of high speed cameras with infra-red sensing capabilities enabled precise measurements of fish behavior in the dark, allowing for the elimination of visual cues from the study of lateral line functionality in rheotaxis.

Some early experiments that have been considered in the past as evidence against the role of the lateral line are not listed in Table S2 due to a lack of a controlled experimental design in the field setting. For example, some species of salmonids, including salmon and trout, were observed to swim against the current in the day and rest on the bottom of a stream at night (***Davidson, 1949***; ***Gibson, 1966***; ***Edmundson et al., 1968***), leading to a conclusion that the lateral line played a minimal role in rheotaxis (***Arnold, 1974***). However, we did not include these experiments for our model validation due to confounding factors posed by the field settings, such as variations in water temperature (***Needham and Jones, 1959***; ***Edmundson et al., 1968***; ***Fraser et al., 1993***) and current speeds at different hours of the day. Daily fluctuations in the availability of food (***Waters, 1962***; ***Elliott, 1965***) is another factor that could influence the activity levels of fish at night, as observed in white bass (***McNaught and Hasler, 1961***) and trout (***Elliott, 1965***). Another class of experiments that led to the previous rejection of lateral line was the demonstration of the imperative role of vision in rheotaxis. Experiments on salmon (***Hoar, 1954***) and herring (***Brawn, 1960***) showed a reduction in rheotaxis when vision was obscured in the dark or in muddy water, conflating the role of visual cues in rheotaxis. Again, these observations do not directly contradict the proposed model, which suggests that in the absence of visual cues, rheotaxis could still manifest provided that the flow speed is sufficiently high.
